# Role of leg campaniform sensilla proprioceptive feedback in *Drosophila melanogaster* adaptive walking

**DOI:** 10.64898/2026.07.22.740025

**Authors:** R.D. Custódio, A. Pierzchlińska, M. Haustein, V. Godesberg, M. Dübbert, T. Bockemühl, E.A. Gorostiza, A. Büschges

**Author notes:** shared senior authorship (E.A. Gorostiza, A. Büschges).

## Abstract

With the newly uncovered access to a genetic line labeling all campaniform sensilla in *Drosophila melanogaster* and the many insights gathered from larger insects, we analyzed the distribution and patterning of this class of proprioceptors throughout the fly nervous system and studied CS function in motor control.

We demonstrate via two-photon calcium imaging microscopy that campaniform sensilla activation induces activity in many leg muscles, showing that these proprioceptive stimuli can influence motor neuron activity. We then dissected how the lack of these proprioceptive stimuli influence walking behavior, leg kinematics, and interleg coordination in freely-walking flies using transient optogenetic inhibition and video tracking with high spatiotemporal resolution. We show that campaniform sensilla inhibition robustly affects *Drosophila melanogaster*’s ability to reach their typical walking speeds. Detailed analysis of leg kinematic suggests that shorter stance amplitudes characterized by their long-lasting duration were behind the speed deficits. Dissecting this phenotype also revealed the inability to maintain coordinated interleg stepping patterns as well as postural control in absence of proprioceptive force and load feedback.

## Introduction

The perception of self-generated action is essential for all behaviors, shaping how animals interact with their environments. This holds for default motor patterns, but more specifically for adapting task-specific motor outputs at situationally relevant moments. This feedback encompasses somatosensation ^1^, including proprioception ^2,3^ – the sense of body and self-movement – and exteroception ^4,5^ – the sense of the body interfacing with the world.

During walking legs cope with two different substrates: the ground and the viscous air ^6^. During stance phases, legs are mechanically coupled to the ground to generate forces for propulsion and posture control ^7,8^. During swing phases, legs are lifted off the ground and, constrained only by mass and passive muscle forces ^6^, move to the starting point of the next stance phase ^9^. Signals regarding leg states are transmitted by proprioceptors to neural circuits in the central nervous system (CNS) generating and coordinating leg movements ^2–4,10^. Two proprioceptive modalities have been shown to play distinct roles for these neural circuits.

Sensory signals about position and movement of the leg and its segments serve the generation of stance and swing phases, as well as the timing of the transitions between them ^2,7,8,10^. In insects, hair plates located on the cuticle close to the joints between leg segments ^11,12^ and chordotonal organs (COs) arranged in parallel to leg muscles ^13,14^ serve this modality together. Dysfunction in *Drosophila* COs ^15,16^ and in mammalian analogues, the muscle spindles ^17,18^, have been shown to cause functional deficiencies during swing phases.

Secondly, load and force signals contribute to execution of the stance phase, during which the fly’s body load is exerted on the leg’s musculoskeletal system and forces are generated by leg muscles. This includes the transitions to and from the swing phase ^8,10^. In vertebrates, Golgi tendon organs monitor force and load ^19,20^, while in insects, these are encoded by cuticular sensors ^21^ called campaniform sensilla (CS) (Diptera ^16,22^; Orthoptera ^23,24^; Blattodea ^7,25,26^). CS are distributed along trochanteral, femoral, tibial, and tarsal leg segments. They are arranged in groups and fields on the trochanter, femur, tibia and tarsus and as single sensors on the femur and tarsus. Together, they comprise dozens of sensory neurons per leg ^7^, specifically, 42 in *Drosophila* ^27^.

In cockroaches, locusts, and phasmids, mechanical stimulation of single CS or CS groups was shown to affect specific motor neuron pools ^7,28,29^. For example, CS sensitive to forces acting on the trochanter were found to assist coxal motor neuron pool activity during stance phase ^30,31^. This highlights task-specific modulation depending on walking direction ^32,33^. Presently, little information exists about contributing sensorimotor pathways. In *Drosophila,* a previous study ^34^ reported 12 bilaterally-projecting CS on the trochanteral field with direct connections to flexor tibiae motor neurons, suggesting that they are involved in stance activity. Similar connectivity between tibial CS and flexor motor neurons was reported in locusts ^35^, suggesting an assisting function for flexor activity during leg movements, including walking.

No conclusive picture existed regarding the processing pathways between leg CS and leg motor neuron pools, nor how their activity impacts walking behavior in freely-walking animals. Therefore, a *Drosophila* line labeling all CS is a critical tool, useful to fill current knowledge gaps and validate previous observations. Such a line ^36^ will allow the community to functionally characterize these somatosensory neurons; as chordotonal organs ^16^ and mechanosensory bristles ^5^ can now be anatomically and physiologically tested – and hair plates are still inaccessible as a whole ^37^ – CS research can now be holistically explored.

## RESULTS

### eyg-Gal4 labels all campaniform sensilla in *Drosophila* legs

We used eyg-Gal4, a genetic line reported in Hopkins et al. 2023 ^36^, to examine how CS arborizations are patterned throughout *Drosophila melanogaster’s* nervous system (Fig. 1A) and how CS mechanosensory inputs may influence walking behavior. Expression of the eyg transcription factor was shown in male legs as exclusive to CS neurons. We verified its expression in female eyg>GFP flies as their size is preferential for detection and annotation of high-speed video recordings for walking behavior assays ^5,15,16^.

**Figure 1:**
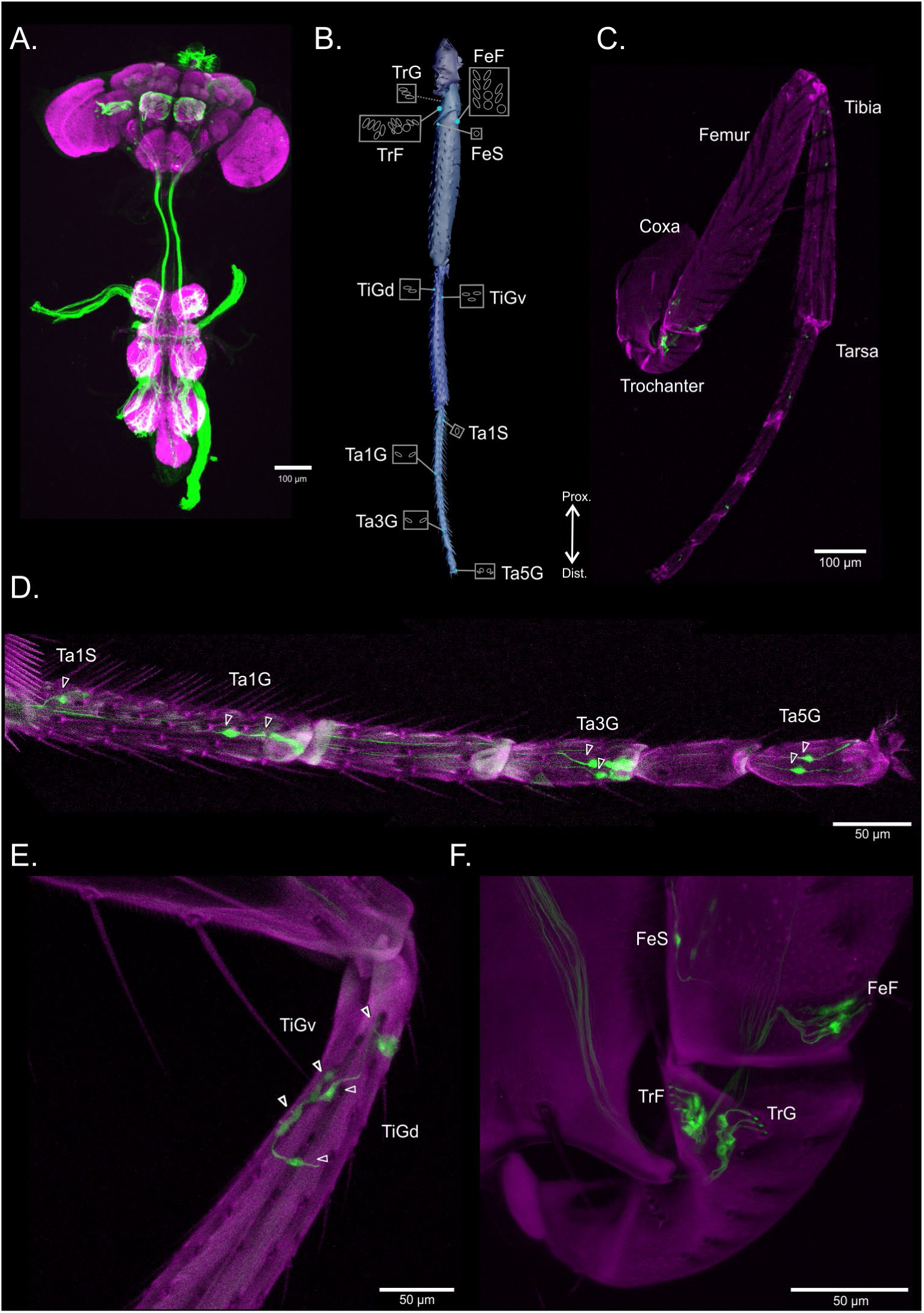
**A)** Max intensity stack of an eyg>GFP *D.mel* CNS (10x) expressing GFP in eyg-positive neurons depicting all CS projections onto the VNC. Magenta: nc82 signal of CNS labeled with Alexa 405 (398-412nm). Green/white (post-processing editing for increased contrast): GFP signal from eyg-Gal4 labeled cells (502-517 nm) (N=3). **B)** Illustration based on ^27^ depicting the distribution of CS throughout the leg. Left to right, top to bottom: TrG: trochanteral group; TrF: trochanteral field; FeF: femoral field; FeS: femoral single CS; TiGv: tibial group (ventral); TiGd: tibial group (dorsal); Ta1S: first tarsal segment, single CS; Ta1G: first tarsal segment, CS group; Ta3G: third tarsal segment, CS group; Ta5G: fifth tarsal segment, CS group. **C)** Max intensity projection of eyg-positive cells in a front leg (10x) of a female eyg>GFP *D.mel* (n=9). Magenta: background autofluorescence of leg cuticle (666-800 nm). Green: GFP signal from eyg-Gal4 labeled cells (502-517 nm), replicating the locations of the SEM-based scheme in Fig. 1B. **D)** Max intensity projection a tile scan of eyg-positive cells in a front leg’s tarsa (40x) of a female eyg>GFP *D.mel* (n=3). Description of channel intervals as in Fig. 1C. **E)** Max intensity projection of eyg-positive cells in a front leg’s tibia and tarsa (40x) of a female eyg>GFP *D.mel* (n=3). Description of channel intervals as in Fig. 1C. **F)** Max intensity projection of eyg-positive cells in a front leg’s coxa, trochanter and tibia (63x) of a female eyg>GFP *D.mel* (n=3). Description of channel intervals as in Fig. 1C.

Confocal fluorescent microscopy of female *Drosophila* legs, wings, halteres and whole fly stereomicroscopy (Fig. 1, Fig. S1) were compared to scanning electron microscopy descriptions of CS numbers and locations on the leg cuticle ^27^. Stereomicroscopy revealed labelling in all known locations (Fig. 1B,C, Fig. S2A, B) ^27^ as well as in the proboscis and antennae (Fig. S1A).

We verified three CS on the first tarsal segment (Ta1S) and two on the third tarsal segment (Ta3G) ^27^. On the fifth tarsal segment (Ta5G) we regularly located two cell bodies (Fig. 1D, Fig. S1C) two less than expected ^27^ but in line with a second study ^38^.

Labeling was harder to differentiate on the tibia therefore we used eyg>GFP,RedStinger in order to individually label eyg-positive somata and differentiate them from other structures. We detected two somata forming the ventral group (TiGv, bold arrows) and three forming the distal group (TiGd) (n=6 legs) (Fig. 1E) ^27^ by co-expressing red (nuclear) and green (membrane) fluorophores co-labeling cell nuclei (yellow) (Fig. S1D).

We observed complex arrangements of eyg-positive neuronal structures in the femur and trochanter which could not be disentangled with eyg>GFP,RedStinger (Fig. S1E). Imaging revealed a lone femoral cell (FeS), the femoral field (FeF), trochanteral group (TrG) as well as a location suiting the two described trochanteral fields (TrFp and TrFa) (Fig. 1F, Fig. S1E) ^27^. These could not be distinguishable into two isolated trochanteral fields due to overlapping fluorescence and were therefore inspected as a whole (TrF).

Despite femoral and trochanteral labeling densities, we confirmed all predicted CS locations and numbers ^27^ by imaging legs from different angles at higher magnifications. We captured one representative image with the optimal alignment (Fig. S2A,B) to visualize all 28 predicted femoral and trochanteral CS (Fig. 1F, Fig. S1E) ^27^.

By selecting specific slice intervals (Fig. S2C-F, Supplementary Video 1) we identified three eyg-positive cells comprising the trochanteral group (TrG) (Fig S2C), twelve comprising the trochanteral field (TrF) (Fig. S2D), twelve comprising the femoral field (FeF) (Fig. S2E) and one lone cell in the femur (FeS) (Fig. S2F). At these magnifications CS show the characteristic cilium-like dendrites emerging from the cuticle, connecting to a somata with an axon ascending through the leg towards the VNC. Therefore we corroborated by number, location ^27^ and morphology that eyg-Gal4 ^36^ labels all leg-CS (Fig. 1C-F, Fig. S1B-E, Fig. S2B-F). Imaging of eyg>GFP flies also revealed labeling in all eight predicted wing locations ^27^ (Fig. S1F) but, due to high cell density ^27,39^, confocal microscopy inspection of the halteres was only verified via stereomicroscopy (Fig. S1A, black arrowhead).

With these findings we concluded that eyg-Gal4 was suitable for a generalized CS optogenetic manipulation within the context of walking behavior.

### eyg-Gal4 activation elicits muscle activity in *Drosophila* leg segments

To study how transient CS loss of function affects walking behavior, it was important to understand which components of the motor system CS interact with. We know that leg CS elicit leg muscle activity in larger insects ^26,31,40,41^ and that some directly connect to fly leg flexor motor neurons ^34^. To understand if CS activity can influence *Drosophila* leg motor neurons we optogenetically activated them using CsChrimson while recording GCaMP6f muscle activity within femoral and tibial leg segments of males and females (Fig. 2, Supplementary Videos 2, 3). We ensured that the registered muscle activity arose from activating a single leg’s CS by amputating the flies’ heads, wings and all legs, except the left front leg. Left front legs were placed in three different configurations – extended, neutral and flexed – with no apparent influence on muscle activation.

**Figure 2:**
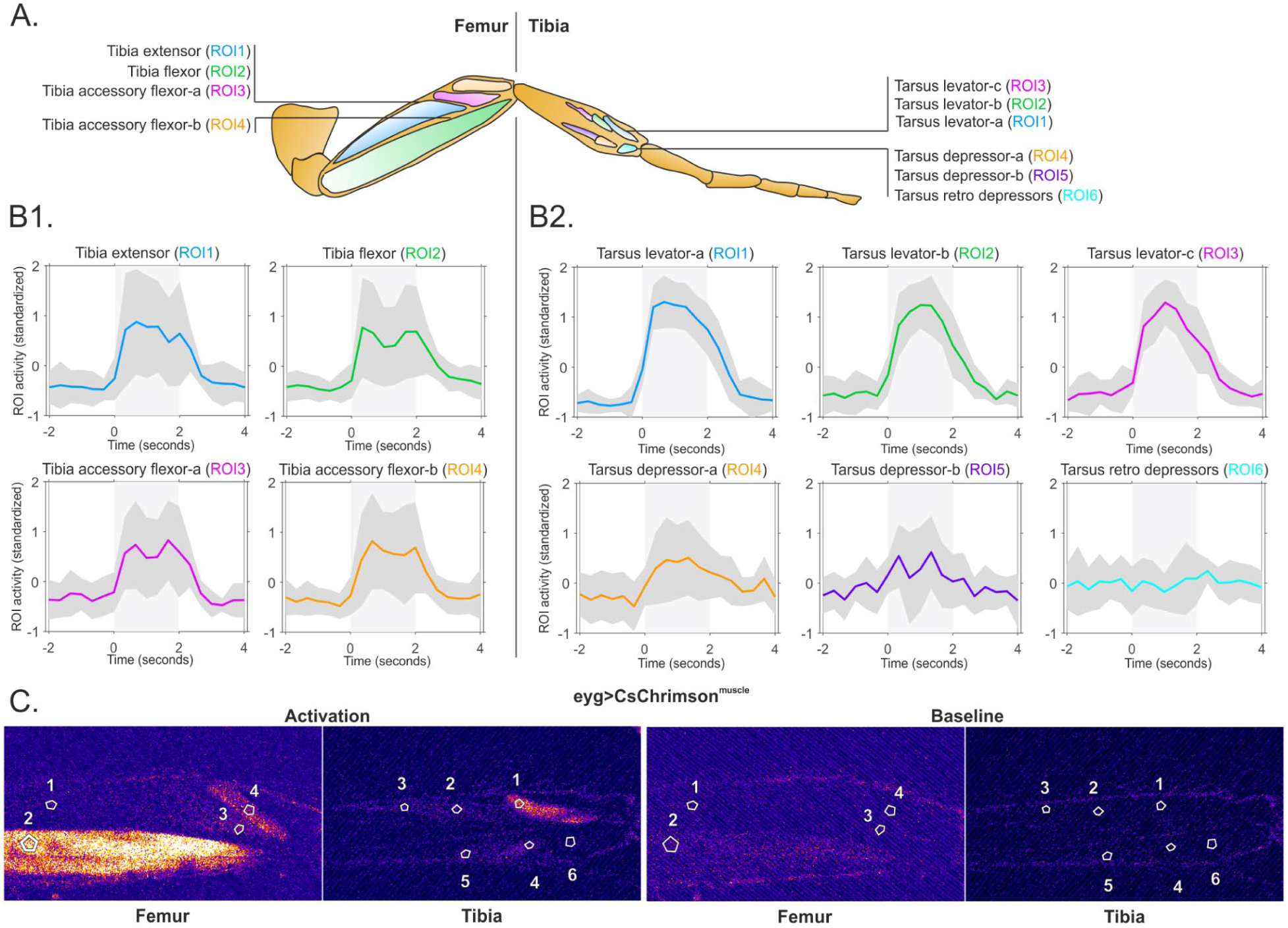
**A)** Depiction of imaged femoral and tibial leg muscles where numbers indicate muscle ROIs in panels B and C. For all muscle nomenclatures please see Azevedo *et al.* 2024 (supplementary materials)^45^ and Soller *et al.* 2004^44^. **B) (B1)** Quantification of femoral muscle activity events in eyg>CsChrimson^muscle^ flies (y-axis: standardized activity within ROI) elicited by a 2 s red light pulse (x-axis: prestimulus (-2 to 0 seconds), stimulus (0 to 2 seconds) and poststimulus (2 to 4 seconds) intervals. ROIs 1 to 4 correspond to leg muscles in graph titles. Colored line depicts mean standardized activity, grey shade depicts SD of activity from different preparations (N=14, pooled data from extended (N=3, n=4), neutral (n=4) and flexed (N=5, n=6) leg positions). **(B2)** Quantification of tibial muscle activity events in eyg>CsChrimson^muscle^ flies (y-axis: standardized activity within ROI) elicited by a 2 s red light pulse (x-axis: prestimulus (-2 to 0 seconds), stimulus (0 to 2 seconds) and poststimulus (2 to 4 seconds) intervals. ROIs 1 to 6 correspond to leg muscles in graph titles. Colored line depicts mean standardized activity, grey shade depicts SD of activity from different preparations (N=12, pooled data from extended (N=3), neutral (N=5) and flexed (N=4) leg positions). **C)** Depiction of a single muscle activity event (left, “Activation”) and baseline muscle activity (right, “Baseline”) within the femoral or tibial leg segments of the same eyg>CsChrimson^muscle^ fly. ROI numbering corresponds to muscles analyzed in (B).

CS activation reliably elicited muscle activity in eyg>CsChrimson^muscle^ femoral and tibial leg muscles. Contrarily, this protocol did not elicit muscle activation in emptySplit>CsChrimson^muscle^ flies (Fig. S3, Supplementary Videos 4, 5), indicating that the light stimulus was not responsible for spontaneous activity. Four femoral regions of interest (ROIs) were selected: tibia extensor, - flexor, -accessory flexor-b and -accessory flexor-a muscles ^42–45^ (Fig. 2A, B1, C). All four were active during a 2 second optogenetic stimulus. Interestingly, tibia flexor and -accessory flexors-a and -b displayed two activity maxima whereas the tibia extensor was continuously active (Fig. 2B1).

Within the tibia we selected six ROIs: three in the tarsus levator fibers ^44,45^ (ROI 1-3), two in the tarsus depressor ^44,45^ fibers (ROI 4-5) and one in the tarsal retro depressor ^45^/ reductor ^44^ fibers (ROI 6) (Fig. 2A, B2, C). We chose to record three ROIs within the tarsal levator as – being large muscle fibers that, to the best of our knowledge, had not yet been recorded – our confidence in the analysis increased with additional ROIs. The same applies for the tarsus depressor muscle. CS activation reliably elicited muscle activation in all tarsal levator muscles. This effect was elicited to a lesser extent in the first ROI of the tarsus depressor. Interestingly no activity was observed in tarsus retro depressors (Fig. 2B2). Concluding, CS signals access motor neurons targeting most femoral and tibial leg muscles.

### Optogenetic inhibition of all eyg positive neurons induce locomotor defects in climbing *Drosophila*

To uncover the role of CS in walking on a systems level we used climbing assays (Fig. 3A) to compare eyg>GtACR1 flies in the dark (internal control) *versus* under GtACR1-induced optogenetic inhibition (Fig. 3A (left), Supplementary Videos 6, 7) (n= 100; the reader is referred to the figure legends regarding the number of animals/ videos for all subsequent experiments). Inhibiting CS in climbing eyg>GtACR1 flies resulted in a 40.9% performance reduction. To exclude influence of light we drove GFP expression replacing the channelrhodopsin GtACR1 (eyg>GFP). Results showed that the light alone did not cause eyg>GtACR1’s climbing performance reduction as eyg>GFP flies’ performance increased by 4.92% during stimulation (Fig. 3A (middle)).

**Figure 3:**
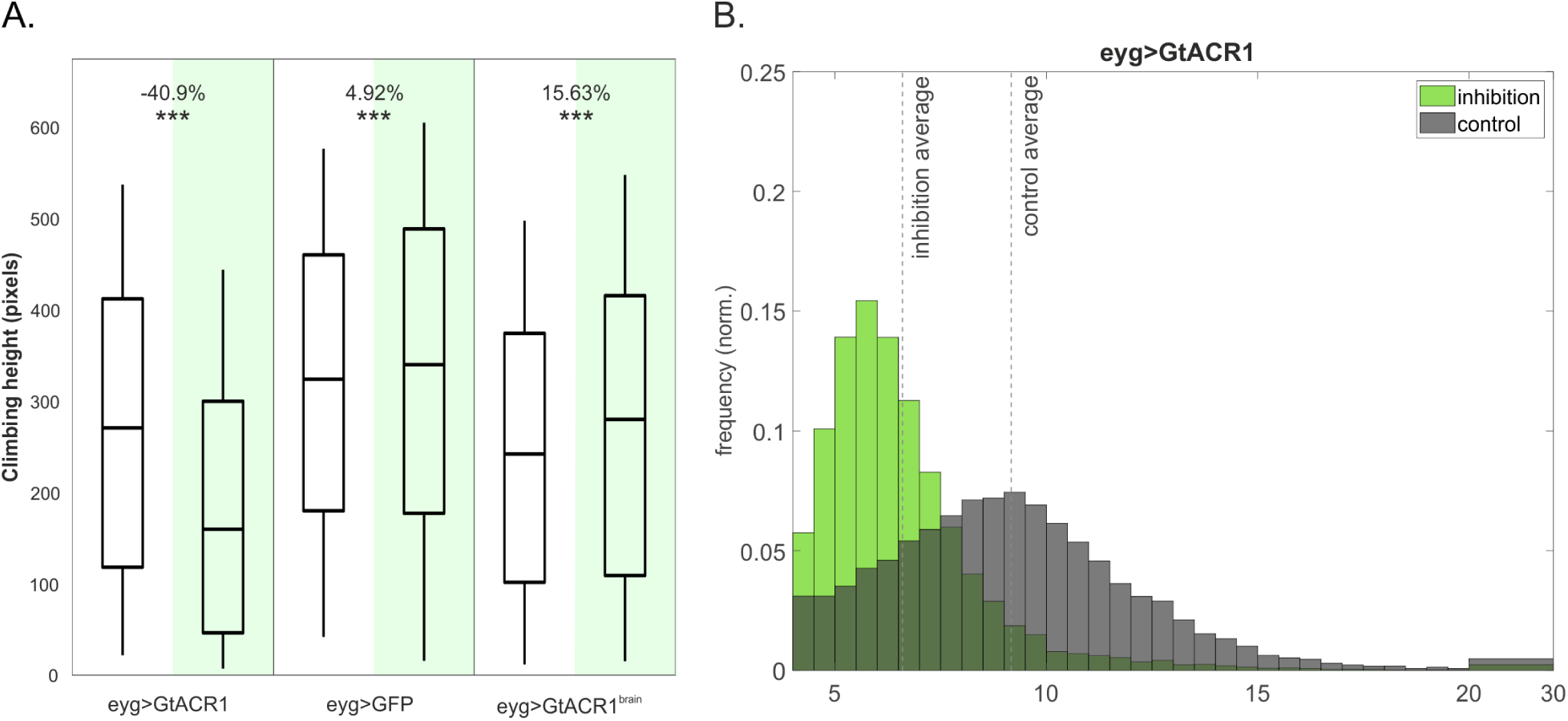
**A)** Assay for negative geotaxis performance (i.e. climbing assay). Comparisons between the experimental group eyg>GtACR1, the brain restriction control eyg>GtACR1^brain^ and the negative genetic control eyg>GFP. White panels display uninhibited climbing performance. Green panels display inhibited climbing performance. The x-axis displays the genotype names. The y-axis displays the climbing height (in pixels); for clarity, see Supplementary videos 6-11. Box and whiskers: the box (top, middle and bottom line respectively) represents the 75, 50 and 25 percentiles of pixel density distribution for the frame for which the climbing height evaluation was performed (42 frames (approx. 2 s) after ground impact). The whiskers represent the upper and lower limits of pixel distribution in the vials (best and worst performers). The ROI for the vials was manually selected using a GUI developed in house. In order to prevent false pixel detection, the top of the ROI was always slightly below the vial plug and the bottom of the ROI was always a bit above the ground (N=10 groups of flies per genotype, 10 trials per group, 100 flies per genotype). **B)** Frequency of speed distribution from walking bouts of control (grey bars) and inhibited (green bars) eyg>GtACR1 flies (N=27). The y-axis represents normalized walking frequency. Vertical dotted lines represent the average of each experimental condition. x-axis represents the walking speed range (mm/s). On the x-axis, the interval between 20 and 30 mm/s pools all the walking tracklets within this speed interval on a single bar for both control and inhibited conditions.

To control for possible effects of the secondary labelling of neuronal structures in nervous system of eyg>GtACR1 flies we investigated this line’s CNS. This showed positive expression of the eyg transcription factor in the proboscis, antennae (Fig. S1A), as well as in brain Mushroom Body (MB), Antennal Lobe (AL) and gnathal ganglion (GNG) neurons (Fig. 1A).

CS presence was never reported on the fly’s head. However, the brain contains central projections from antennal ^46–48^ and mouth apparatus ^43,49^ mechanosensors. It is then conceivable for the eyg developmental transcription factor to be expressed in head mechanosensory appendages. We verified if these neurons contributed to the walking perturbations observed in eyg>GtACR1 flies using the OTD-flippase genetic tool. The eyg>GtACR1^brain^ construct restricted eyg-Gal4 expression to the head and brain ^50^, broadly reducing EYFP expression in eyg>GtACR1^brain^’s legs (Fig. S4A) and VNC (Fig. S4B). Since restriction tools are never 100% effective ^51^ labeling was still observed within VNC neuropils; leg expression was nevertheless considered negligible as the green laser intensity was increased to twice that of Figure 1. eyg>GtACR1^brain^ flies presented no climbing deficits (Fig. 3A (right), Supplementary Videos 8-11), displaying instead a 15.63% increased performance during inhibition.

These experiments indicated that the climbing performance deficits (Fig. 3A (left)) were not caused by silencing eyg-positive neurons in the brain. Therefore, we considered eyg-Gal4 suitable to study CS’s role within the context of *Drosophila* walking behavior.

### Optogenetic inhibition of CS neurons decreases the speed of freely-walking *Drosophila*

The reduction in climbing performance prompted the question on how horizontal walking might be affected without CS feedback. Therefore, we analyzed *Drosophila*’s walking speed during CS inhibition (Fig. 3B) while controlling for genetic background and the influence of labeled brain neurons (Fig. S5A-C).

Transient inhibition in eyg>GtACR1 flies drastically reduced walking speeds (Fig. 3B). While speeds ranged from 4 to 30 mm s^-1^ (with a maximum at 9 mm s^-1^) during control conditions, these decreased to 5 to 15mm s^-1^ (with a maximum at 5.5 mm s^-1^) during CS silencing. Average speed decreased from 9 mm s^-1^ in the internal control, to 6.5 mm s^-1^ upon CS silencing. Importantly, the speed distribution of the internal control group (Fig. 3B) was comparable to that of empty>GtACR1 genetic control flies (Fig. S5C) and to all tested control genotypes (Fig. S5A-B, D-F (18°C), G-I (control)) implying normal walking under unperturbed conditions. Meanwhile, the genetic control eyg>GFP did not show average walking speed differences between the light-on and light-off scenarios (Fig. S5A), confirming that GtACR1 elicited the slow-walking phenotype (Fig. 3A (left), 3B). Again, we confirmed that the effect in walking speed did not relate to neurons labeled in the brain (Fig. 3A (right), Fig S5B).

Walking speed decreases in positive controls were also observed when depleting NompC channels – essential for mechanotransduction ^52,53^ – from CS (Fig. S5E-F). We used the thermogenetic tubulin-Gal80 construct to express NompC interference RNA (RNAi) by exposing flies to a 29°C permissive temperature for three days after hatching (preventing pre-eclosion developmental defects). The experiment was controlled with animals raised at 18°C (restrictive temperature), while all animals were tested at room temperature. Arguably, the fact that flies were reared at different temperatures could create confounding effects since higher temperatures typically promote walking activity ^54,55^ and, conversely, lower temperatures reduce it. However, we showed that eyg,tub^Gal80ts^>NompC^RNAi(TRIP)^ flies walked slower at 29°C than those with intact NompC channels (Fig. S5E (18°C)), confirming that CS mechanosensation deficits were at play in the slow-walking phenotype. This was corroborated using an additional RNAi construct (Fig. S5F). Notably RNAi expression – not different rearing temperatures (Fig. S4D) – promoted the walking speed shift, apparent from the gap in walking speed maxima and average between permissive and restrictive temperatures (Fig. S5 D-E).

By amputating wings and/or halteres ^56^, we also found that CS on these body-parts did not contribute to walking speed (Fig. S5G-I (control)) and that the phenotype arising from CS inhibition was unaltered (Fig. S5G-I (inhibition)). Therefore, we concluded that leg-CS feedback was responsible for the observed walking speed decreases during inhibition.

### Optogenetic inhibition of all eyg positive neurons causes extreme kinematic defects in leg stepping of freely-walking *Drosophila*

In the next sets of experiments we dissected the contribution of CS feedback during walking by measuring detailed leg kinematics of freely-walking flies while transiently inhibiting CS. Comparing straight walking bouts from the same flies during control and inhibition conditions provided an internal control; however drastic speed decreases during CS inhibition prevented speed matching – i.e. comparing control and inhibited walking bouts performed at similar speeds, minimizing its potential kinematic effects.

Our analysis revealed markedly altered walking kinematics during transient CS inhibition in eyg>GtACR1 flies (Supplementary Videos 12, 13). Leg swing durations decreased 14% to 16% in all leg pairs (Fig. 4A1) whereas stance durations increased 72% to 77% (Fig. 4A2), leading to 41% to 46% increased step durations (Fig. 4A3). Importantly, despite stance duration increases, CS inhibition decreased stance amplitudes by 9% to 15% (Fig. 4A4, B1). Furthermore, variability markedly increased in all kinematic parameters but one: swing duration.

**Figure 4:**
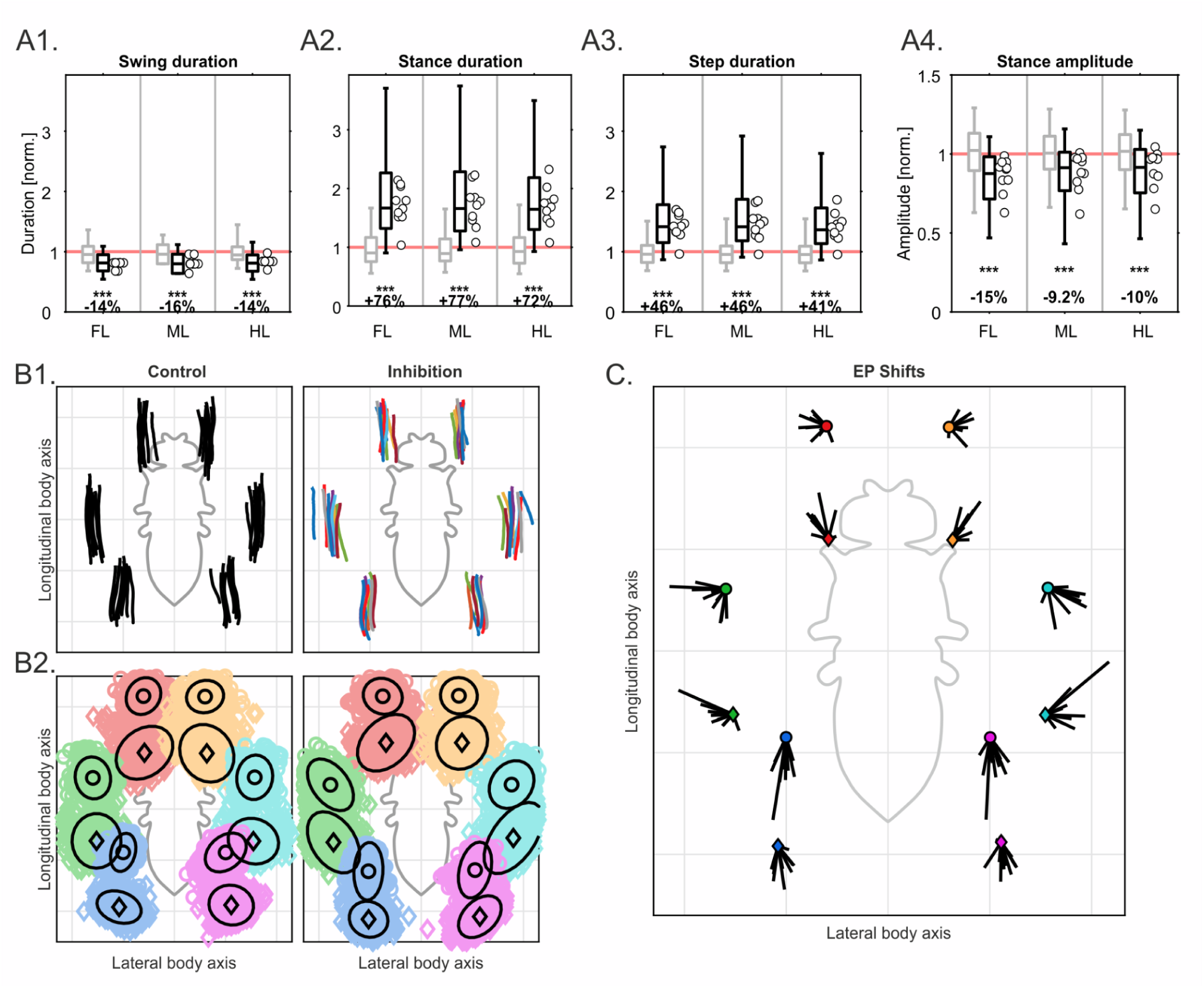
**A)** A1-4) Normalized swing, stance and step durations and stance amplitudes produced during straight walking bouts (minimum 5 steps per leg) by eyg>GtACR1 flies (y-axis). Grey and black boxplots represent walking bouts produced by the same fly in the dark and under green light. White dots: individual average of each inhibited fly. Steps from front (FL), middle (ML) and hind legs (HL) were paired (x-axis). Normalization was done on a per fly basis to their mean values in the control condition (indicated as 1, light red reference line). Box plots (whiskers indicate 1% and 99% percentiles) are combined normalized data for all flies; circles indicate medians for individual flies. Combined data in the inhibited condition were tested against a median of 1 with a two-sided Wilcoxon signed rank test. Significance levels of these tests are given as non-significant (n.s.), *P≤0.05, **P≤0.01 and ***P≤0.001. Effect sizes are given as the difference between the median in the inhibited condition and 1 (normalized control condition). Each fly (N=9 has 10 control and 10 inhibition videos (n=190). **B) B1)** Average stance trajectories for each individual fly shown in fly-centered coordinate space for the control (left: black lines) and inhibited condition (right: lines color coded per individual). **B2)** distribution of anterior (AEPs: colored circles) and posterior extreme positions (PEPs: colored diamonds) with their mean positions (black circles and diamonds) and 95% percentiles (black ellipses). Colors represent the front (right: red; left: yellow), middle (right: green; left: cyan) and hind legs (right: blue; left: pink). **C)** Extreme position (EP) shifts. For each leg, each vector (black lines) corresponds to one fly and displays the average shift of AEPs (top) and PEPs (bottom) in the inhibited condition, compared with their respective mean positions in the control condition (circles, color coded as B2.).

Additionally, we observed alterations in the spatial control of leg placement during walking. Anterior extreme positions (AEPs) and posterior extreme positions (PEPs) displayed increased variability – evident from wider, elongated ellipses (Fig. 4B2) – and a broader walking posture (Fig. 4C; black lines *versus* colored circles and diamonds). The same analysis was performed in eyg>GtACR1^brain^ flies, conclusively demonstrating that brain and head mechanosensitive neurons did not contribute to the CS silencing kinematic phenotype (Fig. S6, Supplementary Videos 14, 15). Since – even at the level of detail provided by high spatiotemporal leg kinematic analysis – eyg>GtACR1^brain^ flies displayed no locomotor phenotype, we concluded that the slow-walking speeds stemmed from inhibited leg, haltere or wing CS networks. As no speed alterations occurred in absence of wings and/or halteres (Fig. S5G-I) the underlying contributor to altered stepping was the absence of leg-CS feedback.

### CS inhibition disturbs interleg coordination in straight walking

Reduced walking speeds (Fig. 3B) and increased stance durations coupled with decreased amplitudes (Fig. 4A2,A4) raised the hypothesis of disturbed interleg coordination during CS inhibition. Flies perform interleg coordination patterns dependent on walking speeds ^15,57^, typically adopting the tripod coordination pattern at medium to high walking speeds. Independent of speed, alternating and non-overlapping legs swing in a hind-to-front fashion – i.e., anterograde swing progression ^58,59^ – resulting in the same number of stepping cycles across legs. This situation occurred for control eyg>GtACR1 (Fig. 5A1, Supplementary Video 13) and for control and inhibited eyg>GtACR1^brain^ flies (Fig. S7A1-2, Supplementary Videos 14, 15). However, upon CS silencing, these interleg coordination patterns deteriorated (Fig. 5A2-3, Supplementary Video 12): step cycle counts between legs differed, the back-to-front sequencing of ipsilateral legs stepping and the alternation between contralateral legs were no longer displayed systematically. Individual legs generated two successive swing phases without intermittent contralateral leg swings and individual legs skipped the anterograde swing progression. Additionally, due to decreased walking speeds, eyg>GtACR1 inhibited flies needed more time to perform approximately the same number of steps as compared to control (Fig. 5A1-3).

**Figure 5:**
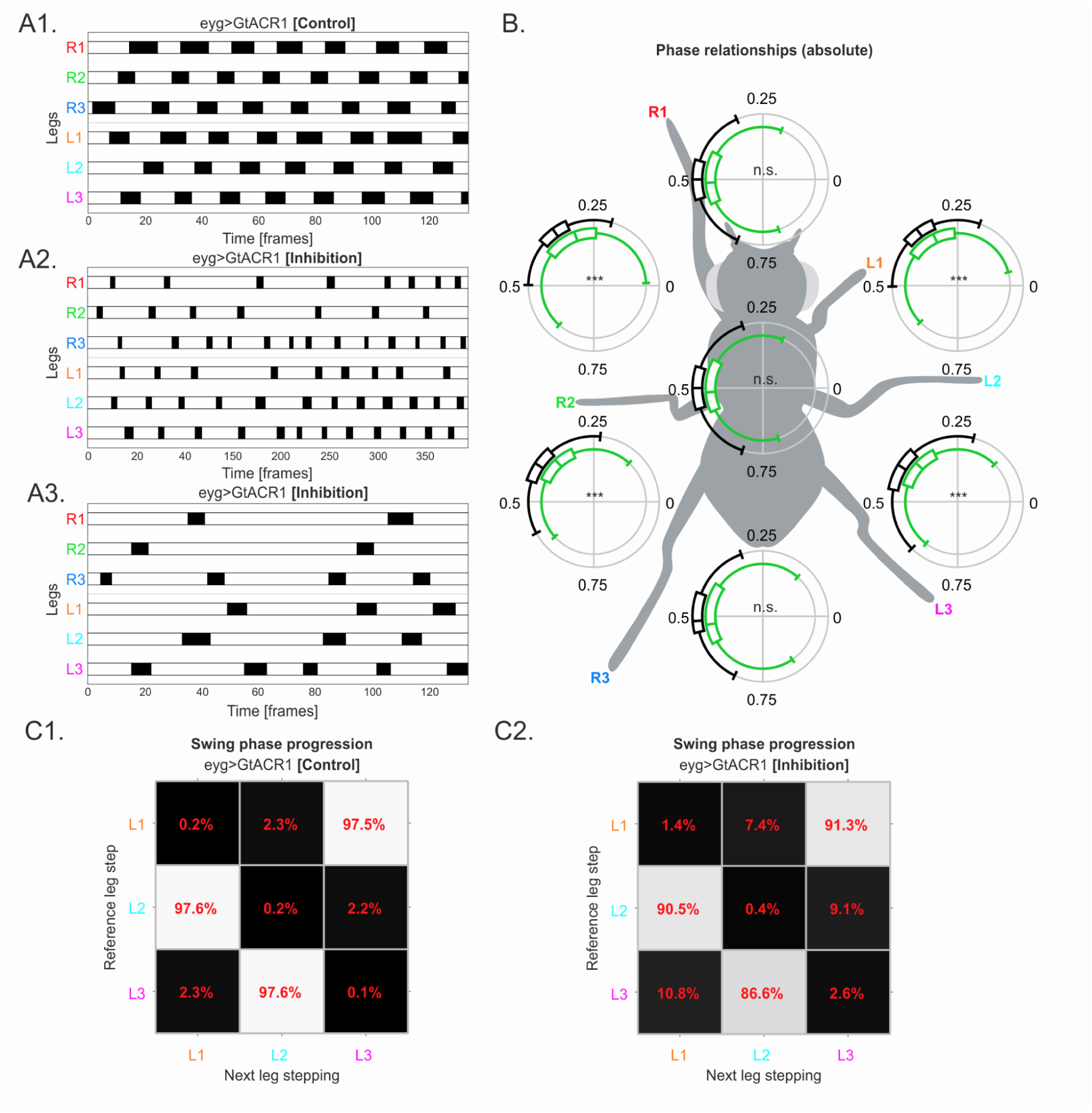
**A)** A1) Footfall patterns of example trial from a eyg>GtACR1 fly walking in the dark (control) and **A2)** light (inhibition) performing a straight walking bout covering approximately the same distance. The x-axis represents time in frames. In the y-axis R1, R2, R3 and L1, L2, L3 respectively account for the right and left front, middle and hind legs. White and black bars respectively represent the legs stance and swing phases. **A3)** Snippet of A2) with the same number of frames as A1) (frame interval: 150-283). **B)** Comparison of absolute leg phase relationships. Circular boxplots placed between corresponding leg pairs for eyg>GtACR1 flies. On each circle, black boxplots are uninhibited control walking bouts and green boxplots are inhibited walking bouts depicting interleg coordination. Combined data in the inhibited condition were tested against the data in the control condition with a non-parametric variant of the Kruskal–Wallis test for circular data. Effect sizes are given as absolute differences between the median of the inhibited condition and 0. Absolute effect sizes smaller than 0.005 have been rounded to 0. Exemplifying with R3 and L3 phase relationships, control (black) depicts the median antiphasic relationship at 0.5. Inhibition (green) depicts the median as in antiphase stepping (0.5); 75 and 25 percentiles almost reach phasic stepping (0), i.e. legs swinging or standing at the same time. Walking bouts are the same from Fig. 6. Each fly (N=9) has 10 control and 10 inhibition videos (n=190). **C)** Depiction of back-to-front ^58,59^ swing phase progression between ipsilateral left legs for uninhibited (control: top matrix) and inhibited (bottom matrix) walking bouts in eyg>GtACR1 flies. Reference legs are depicted on the y-axis, next stepping leg is depicted on the x-axis. Percentiles on squares show the probability of a specific leg stepping. Each fly (N=9) has 10 control and 10 inhibition videos (n=190).

Quantification of phase relationships between stepping legs revealed alterations in interleg coordination (Fig. 5B). For contralateral leg pairs – R1-L1, R2-L2 and R3-L3 – the median phase value comparisons between control and CS inhibited conditions were statistically insignificant, however variability during inhibition largely increased, i.e. whiskers beyond 0.75 and below 0.25.

For ipsilateral legs – R1-R2-R3 and L1-L2,L3 – control trials displayed back-to-front sequencing with average phase values slightly deviating from perfectly antiphasic – i.e. <0.5 – due to phase lag between ipsilateral stepping legs ^15,57,58,60^. CS inhibition remarkably altered interleg coordination. For example, swings of neighboring legs happened close in time – indicated by green whiskers reaching close to “0”, e.g. for the pairs R1-R2 and L1-L2 – an unusual phenomenon as ipsilateral legs normally generate systematic swing alternations across speed ranges ^15,57–59^ (e.g. R1-R2, Fig. 5B).

Similarly back-to-front swing phase progression between ipsilateral legs ^5,15,57–59,61^ was affected as individual legs skipped swings, e.g. for the 2^nd^, 3^rd^, 5^th^ and 6^th^ presumed sequence starting with R3 in Fig. 5A2. This deviation from control became apparent when comparing the probability of ipsilateral leg swing progression in control and inhibited CS conditions (Fig. 5C1-2). Taking L3 as reference, L2 swung after it 97.6% of times, as did L1 after L2 (Fig. 5C1). In absence of CS feedback the likelihood of L2 stepping after L3 reduced to 86.6%, L1 after L2 to 90.5% and L3 after L1 to 91.3% (Fig. 5C2). This means that at least one out of 10 steps were not coordinated, differing markedly from control, occurring one out of 50 steps. Expectedly, eyg>GtACR1^brain^ flies displayed no effects on interleg incoordination (Fig. S7).

Given these findings we conclude that CS sensory feedback is crucial for interleg communication and coordination during walking.

### Optogenetic inhibition of all eyg positive neurons causes postural changes in freely-walking, but not standing *Drosophila*

Lastly we tested whether transient CS inhibition affected posture in walking and standing flies by recording eyg>GtACR1 from the side through a transparent corridor. These experiments were controlled using eyg>GFP flies as these provided a solid genetic control (Fig. 3A, Fig. S5A).

First hints for CS silencing affecting posture emerged from wider sprawled AEPs and PEPs when walking (Fig. 4C). Ground clearance in eyg>GtACR1 inhibited walking flies dropped, shown by synchronous and persistent decreases in average notum height (Fig. 6A1, B1 (light onset), Supplementary Video 16). This was corroborated in the violin plots, for which the 25, 50 and 75 percentiles shifted below the pre-stimulation baseline height at stimulation onset (Fig. 6B1 (light onset)). Conversely, ground clearance increased upon termination of inhibition, shown by the average ground clearance and violin plot distribution (Fig. 6B1 (light offset), Supplementary Video 17). eyg>GFP control flies were unaffected by light or its absence (Fig. 6C1, Supplementary Videos 18, 19), indicating that reduced ground clearances were not because of eyg>GtACR1’s genetic background.

**Figure 6:**
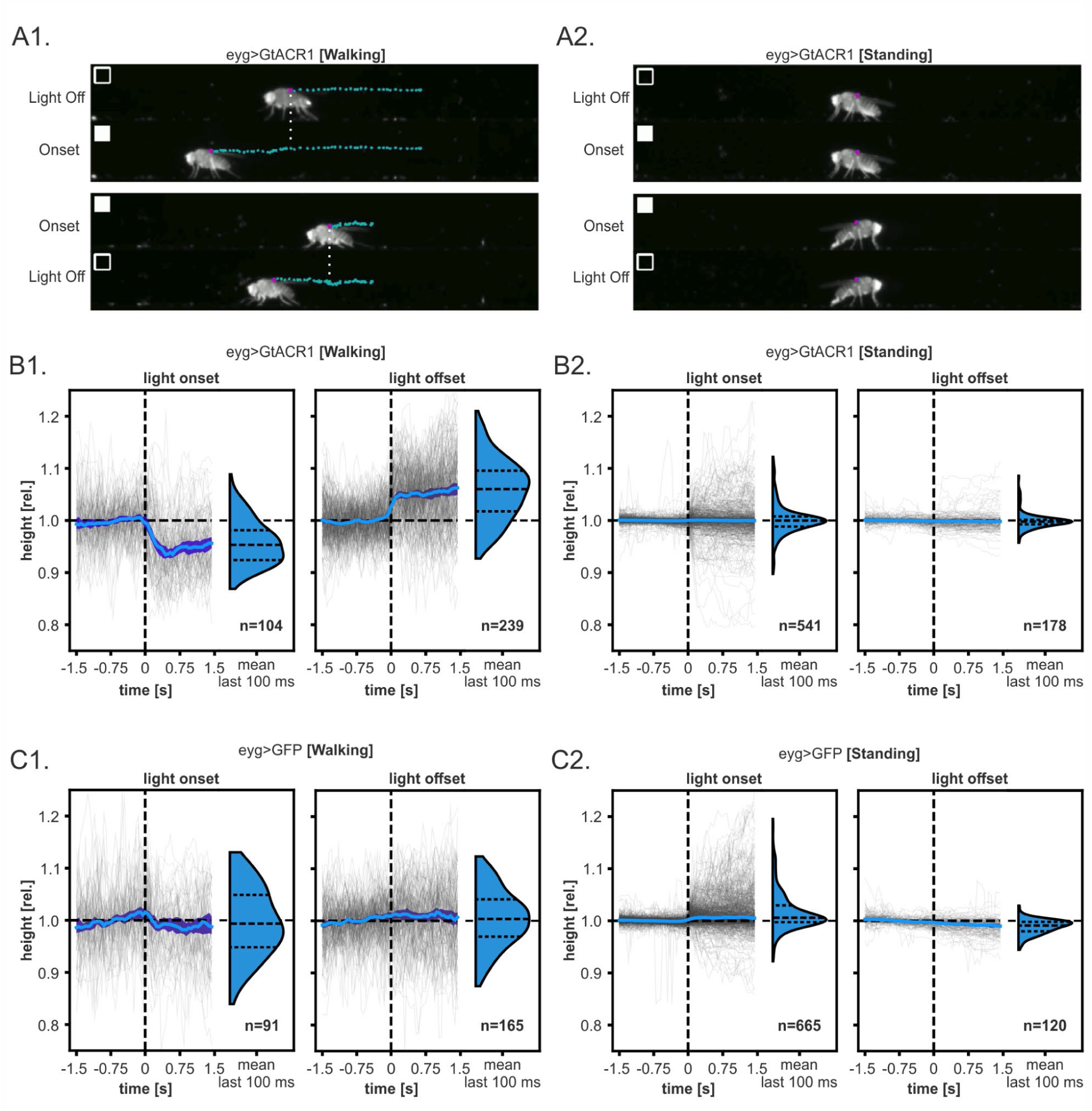
**A)** Exemplary frames of effects analyzed in B). **A1) Top:** “Light off” row shows walking eyg>GtACR1 fly’s ground clearance (height of fly’s notum) before optogenetic inhibition. “Onset” row shows walking eyg>GtACR1 fly’s ground clearance during optogenetic inhibition. Blue dotted traces represent height, frame by frame (dot by dot). White dotted line represents the moment when light turns on (Top) and off (Bottom). **Bottom:** “Onset” row shows walking eyg>GtACR1 fly’s posture at the last frame of CS optogenetic inhibition. “Light off” row shows the fly’s ground clearance after CS optogenetic inhibition is terminated, reflecting a return to baseline ground clearance. “Light off” frame is the last frame of the video. **A2) Top and bottom:** “Light Off”, “Onset” and “Light On” frames represent the same timepoints as A1 in the same eyg>GtACR1 fly standing still instead of while walking. Description of graphic features in Fig. 6A1 legend. **B) B1)** Ground clearance changes for walking eyg>GtACR1 flies (N=13<14; number of flies is between 13 and 14, as we could not collect onset or offset trials from all flies) upon CS optogenetic inhibition light onset (left) and offset (right). The x-axis describes the relative height for which “1.0” is the fly’s baseline walking height. The y-axis displays time (seconds), before and after light onset (left) and offset (right), as depicted by the dotted vertical line. The individual walking trials are represented by the grey traces aligned to the “0” timepoint. Blue lines show the relative median height and dark blue areas represent the 95% CI. Violin plots show the distribution of the mean relative heights of the last 100 ms. **B2)** Representation of height changes of immobile, i.e. standing eyg>GtACR1 flies (N=13<14) during CS optogenetic inhibition onset (left) and offset (right). **C) C1)** Ground clearance changes for walking eyg>GFP genetic control flies (N=15<16). **C2)** Representation of height changes for immobile, i.e. standing eyg>GFP flies (N=15<16) during CS optogenetic inhibition onset (left) and offset (right). Description of graphic features in Fig. 6B1-2 legend.

Interestingly, performing the same experiment on immobile, i.e. standing flies, showed no alterations in ground clearance in eyg>GtACR1 or eyg>GFP flies (Fig. 6A2, B2, C2, Supplementary Videos 20-23). This revealed an interesting conclusion: while CS load feedback has relevance for posture control in walking flies, its transient cessation is apparently no challenge for keeping body posture while standing.

## Discussion

In this study we provided first evidence of how force and load proprioceptive signals conveyed by campaniform sensilla contribute to walking behavior in an insect. In general, insect CS constitute a sensory system functionally similar to vertebrate Golgi tendon organs ^20^. They offer the opportunity to investigate how load and force feedback contribute to complex behaviors in experimentally more accessible animals. Here, we used *Drosophila melanogaster*’s genetic toolkit to access and optogenetically influence its CS during motor output. This was facilitated by a previously reported GAL4 line ^36^ which we found to label all CS, including all 42 in the legs ^27^. While transient CS activation induced broad leg muscle activation, inhibition impaired behavioral performance in different paradigms (climbing and level walking) at distinct levels (speed, posture, leg kinematics, and interleg coordination).

CS optogenetic activation induced activity in most femoral and tibial *Drosophila* leg muscles (Fig. 2), including tibial flexor and extensors ^5,42,44,45^, as well as tarsus levator and depressor muscles ^44,45^. Excitatory connections between leg CS and tibial flexor motor neurons were previously shown ^34^ and, based on findings in other insects, probably exist for other CS. Studies using CS mechanical stimulation reported activation of tibial flexor and extensor muscles ^26,40,41,62^. Due to preparational restrictions we did not explicitly monitor activity in proximal muscles; however, it is likely that CS similarly excite them in *Drosophila;* studies in other insects show that trochanteral CS excite coxal retractor and protractor muscles, as well as trochanteral depressors ^63–66^. Thus, our findings complement these studies’ conclusions that CS feedback acts as a general excitatory source for leg muscle activity. It is however unlikely that this influence is purely excitatory. Studies showed that CS can have inhibitory effects on leg muscles at rest and during active movements ^33,67^. These inhibitory influences are much weaker, probably also in *Drosophila*; since we activated all CS, the higher strength of excitation putatively dominated the observed effects, potentially masking weaker inhibitory influences. To differentiate excitatory from inhibitory influences the field will require a detailed mapping between specific CS and muscles, along with similar investigations using sparser labeling lines.

Silencing all CS during climbing (Fig. 3A) and level walking (Fig. 3B, Fig. 4-6) further supported the notion of an important role for CS in *Drosophila* walking behavior. Without CS feedback, flies showed a distinctly reduced climbing ability (Fig. 3A). Furthermore, they walked much slower and hardly reached or maintained walking speeds above 10 mm s^-1^ (Fig. 3B), a speed that unperturbed animals frequently exceed by a large margin ^15,43^ (Fig S5 (controls)). Similar reductions in walking speed have been reported when all exteroceptors and proprioceptors, including CS, were inactivated ^15^. Interestingly, no similar reduction was observed when COs ^16^ or hair plates ^37^ were inactivated. This points towards a particular role for CS activity in achieving certain walking speeds.

It is known that functional execution of leg movements in walking vertebrates and invertebrates is dependent on movement, force and load signals ^3,4,8,68,69^. Load signals are particularly important for force generation during stance as shown in reduced invertebrate ^30,32^ and vertebrate ^70^ preparations. These also showed that legs will lift off, swinging when their own load decreases, due to changes in other legs’ states. We corroborated and extended these conclusions to intact, freely-walking flies exclusively lacking CS feedback. This resulted in stepping kinematic alterations that robustly affected all legs, characterized by significant stance amplitude decreases and simultaneous stance duration increases (Fig. 4A) as body load redistribution is not sensed. These findings underline the pre-established role for load signals, especially during stance phases.

These effects clearly differed from those reported when other proprioceptors such as leg COs are transiently silenced in *Drosophila* ^16^. Stance and swing durations as well as stance amplitudes were affected; crucially, these effects are leg-specific and their magnitudes increase (stance duration and amplitude) or decrease (swing duration). In contrast, the effect sizes of CS silencing shown here were largely independent of specific legs. We therefore hypothesize that CS provide more generic and global information, while movement information is processed more locally and geared towards the kinematic properties of individual legs. Two additional effects of CS silencing deserve attention: the absence of CS signals made the liftoff and touch-down positions more variable while posture became more sprawled. Furthermore, during CS silencing swing durations decreased by approx. 15%. The latter effect is particularly unexpected: typically, swing durations slightly increase for all legs at slower walking speeds ^57^. We observed the opposite as the animal almost always lost balance and needed more ground support. This could be explained by either a more direct correlation between stance amplitude – decreased here – and swing duration, or, more hypothetically, a role of load information not only in the transition from stance to swing, but also in the maintenance of swing phase. In summary, our results suggest that spatial and temporal aspects of leg stepping during walking are under continuous influence of load signals.

In addition to single-leg kinematics, we also observed clear changes in interleg coordination. Without CS information the variability of phase relationships between adjacent legs strongly increased (Fig. 5B). Often, interleg coordination did not fall into commonly observed patterns like wave, tetrapod, or tripod coordination ^15,57^ (Fig. 5A2-A3) – speed-dependent in walking insects. Wave gait is performed at low walking speeds, tetrapod-like coordination at low to intermediate speeds, and tripod coordination at intermediate to high walking speeds. Independent of the coordination pattern swing phases do not overlap and ipsilateral legs swing in a fixed anterograde directed sequence; this latter aspect is especially strict in wild types and our control conditions and was, with rare exceptions, invariant (Fig. 5C1, Fig. S7C) ^57,71–73^. Here, we regularly observed simultaneous leg-swings executed by adjacent legs (Fig. 5B) and deviations from the anteriorly-directed swing phase progression (Fig. 5C2), strongly indicating interleg coordination defects. This suggests that CS load information plays a role in maintaining interleg coordination, possibly via temporal effects conveyed by body weight redistribution between legs, providing proper phasing; in line with preliminary data demonstrating direct CS inputs onto intersegmental coupling neurons ^74^. Furthermore, interleg coordination without CS feedback seems, to some extent, to replicate effects observed when flies carry additional load ^61^. The two findings seem contradictory but might be reconciled by assuming that in both cases the effective dynamic range signaled by CS is, for all purposes, either strongly restricted (load-carrying case, in the context of Weber’s law) or set to zero (CS inhibition, this study). Both manipulations would effectively remove dynamic aspects of CS signaling, in line with the notion that dynamic aspects of load perception are relevant for interleg communication in *Drosophila*.

One particular aspect of postural changes during CS inhibition is noteworthy: when we investigated ground clearance (Fig. 6) we found that its reduction only happened when flies actively moved, i.e. during walking (Fig. 6B1). When they were stationary, inhibition did not seem to have a direct effect (Fig. 6B2). This might simply have to do with the number of legs that support the fly in each condition: during walking, some legs are in swing at any given time; support is reduced and, in combination with CS inhibition, this might lead to lower force development and lower ground clearance. A more interesting hypothesis might be that the current state of the animal (walking vs. stationary) influences the way CS signals are processed. In this notion, postural control and leg kinematics would only be affected if the animal switches into the walking state during which CS signals fluctuate and are considered for control. During inactivity, the postural control system would switch to a more centrally dominated mode during which load signals are stable and disregarded.

How do these effects compare to those of mammalian proprioceptive loss for locomotor control? Few studies addressed loss of function of leg proprioception. Presently, comparable data is available regarding silencing muscle-spindles, i.e. leg movement feedback, in mice. In Egr3 mutants, loss proprioceptive limb movement feedback was found to cause swing duration alterations ^17^ – corroborating COs’ role in *Drosophila* walking. Contrary to CO loss of function, this mutation did not affect stance durations ^18^. No data are presently available on the effect of silencing force and load feedback from the Golgi-tendon organs in mammals. This highlights the need for parallel research across the evolutionary tree to dissect the modus operandi of such a ubiquitous sensory system, in light of its functional relevance for each species.

In summary, we used inhibition of all campaniform sensilla in *Drosophila melanogaster* to explore the role of load and force feedback in the context of walking. Our results indicate that this sensory modality plays an important role in walking, both for single leg stepping kinematics and interleg coordination. We show that even in very light-weight animals with a body mass of approximately 1 mg, load information and the specific timing of forces are necessary to produce functional motor output for walking. Crucially, we have studied this in intact, freely-walking animals; previous studies used reduced preparations or strong experimental constraints. Here, we manipulated the complete CS complement at once. Future studies will need to focus on the contribution and connectivity of individual or small CS subsets. Clearly, the distribution, arrangement, and orientation of CS on *Drosophila’s* legs is very structured and suggestive of functional specialization; investigating this will further our understanding on the role of load sensing in the context of walking.

## Materials and methods

### Fly rearing

Flies were crossed and reared on standard food ^75^ on a 12h:12h light:dark cycle. They were housed at 25°C, 50-70% humidity, except for temperature sensitive genotypes (Fig. S5D-F) which were reared at 18°C, 50-70% humidity, and, after eclosion, kept at 18°C or 29°C, 50-70% humidity, for three days before experiments. All flies were tested at room temperature (approximately 22°C). For optogenetic experiments, flies were kept on food containing all-trans retinal (ATR, 42 µL of a 100 mmol l^-1^ solution), occluded from light for three days, starting at one to two days post-eclosion. Flies were three to five days old during experiments. The tested genotypes were obtained via the following crosses:

● eyg>GFP i.e.;;eyg-Gal4/UAS-GFP:;;eyg-Gal4/TM6b x;;UAS-GFP
● eyg>GFP,RedStinger i.e. UAS-RedStinger/+;;eyg-Gal4/UAS-GFP:;;eyg-Gal4 x UAS-RedStinger;;UAS-GFP
● eyg>CsChrimson^muscle^ i.e.;UAS-CsChrimson/LexAop-GCaMP6f;eyg-Gal4/MHC-LexA:;UAS-CsChrimson;eyg-Gal4 x;LexAop-GCaMP6f;MHC-LexA
● emptySplit>CsChrimson^muscle^ i.e. UAS-CsChrimson/+;emptySplitAD-Gal4/LexAop-GCaMP6f;emptySplitDBD-Gal4/MHC-LexA:UAS-CsChrimson;emptySplitAD-Gal4;emptySplitDBD-Gal4 x;LexAop-GCaMP6f;MHC-LexA
● eyg>GtACR1 i.e.;;eyg-Gal4/UAS-GtACR1:;;eyg-Gal4/TM6b x;;UAS-GtACR1
● eyg>GtACR1^brain^ i.e. UAS-stop-GtACR1/+;Otd-FLP/+;eyg-Gal4/+:;;eyg-Gal4/TM6b x UAS-stop-GtACR1;Otd-FLP/CyO
● empty>GtACR1 i.e.;;empty-Gal4/UAS-GtACR1 :;;empty-Gal4 x;;UAS-GtACR1
● eyg,tub^Gal80ts^>GFP i.e. tubGal80ts;;eyg-Gal4/UAS-GFP: tub-Gal80^ts^/+;;eyg-Gal4/TM6b x;;UAS-GFP
● eyg,tub^Gal80ts^>NompC^RNAi(TRIP)^ i.e. tub-Gal80^ts^/+;;eyg-Gal4/UAS-NompC-RNAi(TRIP): tub-Gal80^ts^,;;eyg-Gal4/TM6b x tub-Gal80^ts^;;UAS-NompC-RNAi(TRIP)
● eyg,tub^Gal80ts^>NompC^RNAi(val1)^ i.e. tub-Gal80^ts^/+;;eyg-Gal4/UAS-NompC-RNAi(val1):;;eyg-Gal4/TM6b x tub-Gal80^ts^;;UAS-NompC-RNAi(val1)

### Fluorescence microscopy

Confocal images were acquired using a SP8 confocal microscope (Leica, Wetzlar, Germany, RRID: SCR_018169). 10x air or 20x glycerol immersion objectives were used to image the central nervous system (CNS), i.e. brains and ventral nerve cords (VNCs). 10x air, 40x oil, or 63x water objectives were used to image legs and wings. Wavelengths were of 405, 488, and 638 nm. Z-stacks were acquired with a minimum of 100 steps. Maximum intensity projections were created using Fiji. Channel intervals for background autofluorescence of leg cuticles were acquired within 666-800nm (magenta/cyan), GFP signal from eyg-Gal4 labeled cells within 502-517nm (green) and dsRed signal labeling the nuclei of eyg positive cells (UAS-RedStinger) within 575-590nm (red). For Figure 1 and Supplementary Figures 1 and 4 the acquisition resolution was of 1024x1024px and 2048x2048px for Supplementary Figure 2.

### Immunohistochemistry for VNC and brains

Dissections were performed on three to eight day-old adult female flies. Flies were anesthetized with CO_2 a_nd briefly soaked in 70% alcohol to dewax the cuticle. CNSs and legs were dissected in 0.1 mol l^-1^ phosphate-buffered saline (PBS). Samples were fixed in 2% paraformaldehyde (PFA) with glacial acetic acid for 45 min and washed with PBS. The blocking solution contained 5% normal goat serum (NGS) in 0.5% Triton X-100 in 0.1 mol l^-1^ PBS (0.5% PBT). After 30 min of blocking at room temperature, CNSs were incubated with primary antibodies (mouse anti-nc82, 1:25; chicken anti-GFP, 1:500, 1% NGS in 0.5% PBT; Table 1) overnight at 4°C. Finally, CNSs were washed with 0.5% PBT, mounted in VECTASHIELD and covered with a coverslip.

**TABLE 1:**
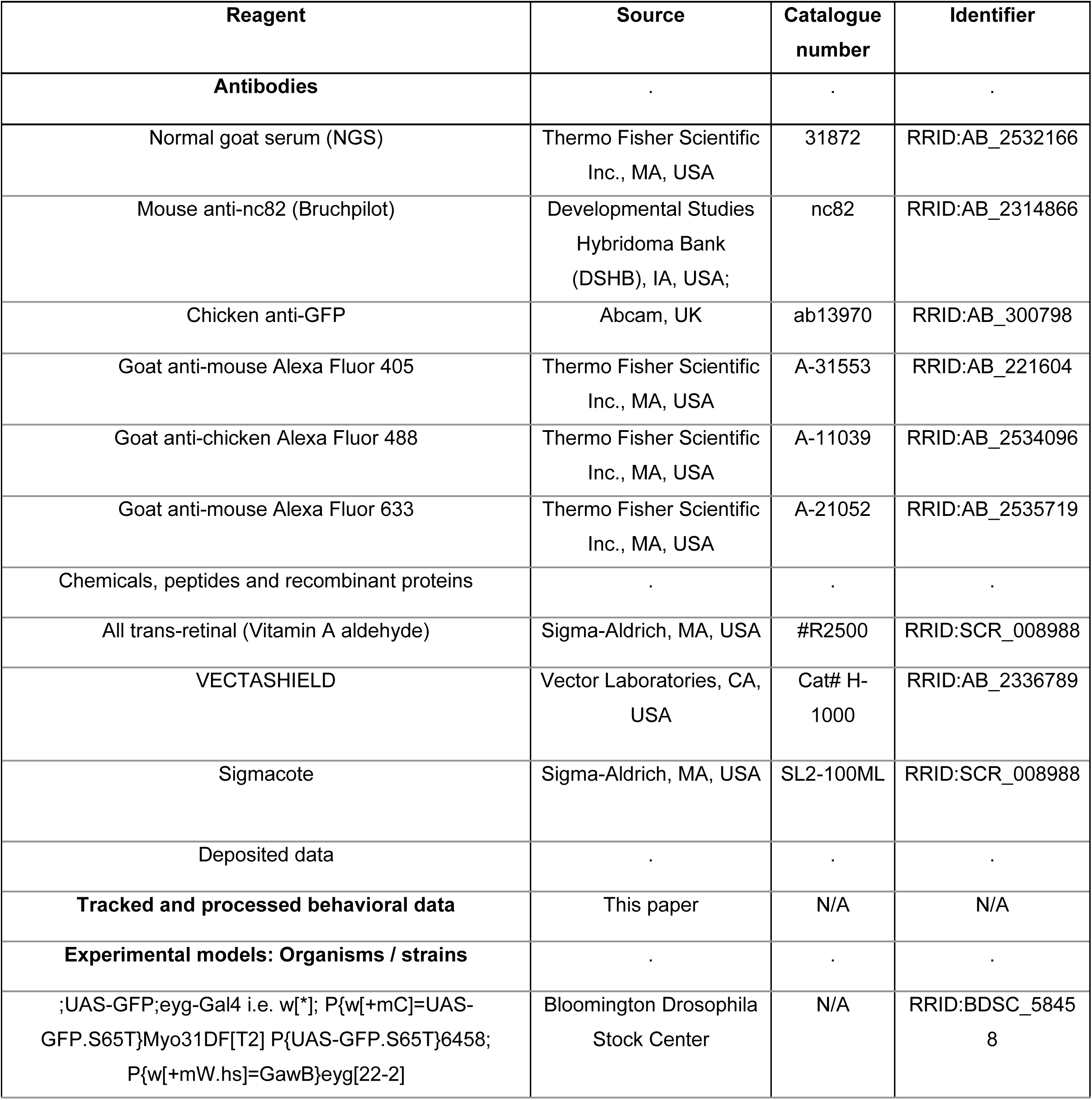
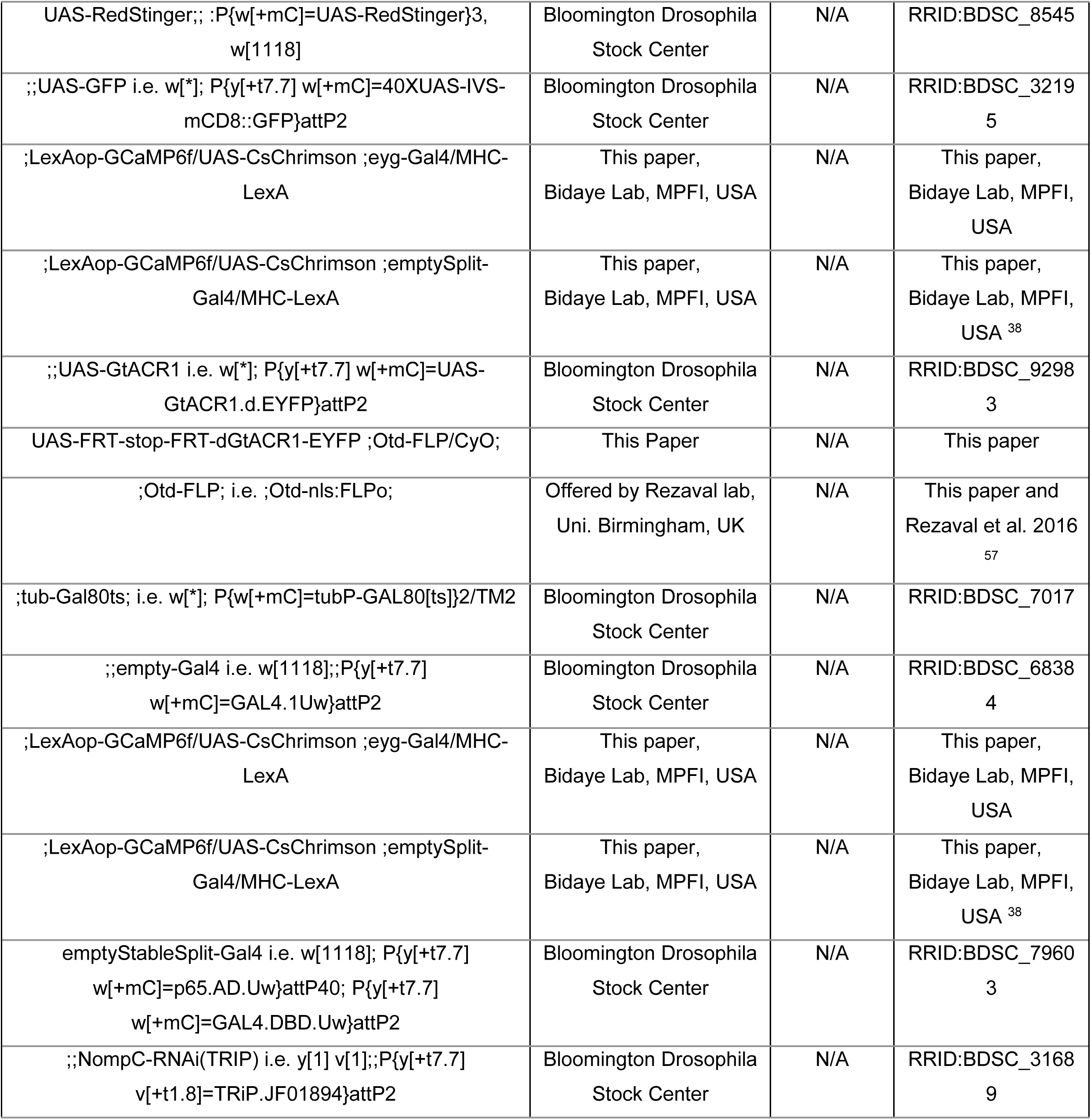
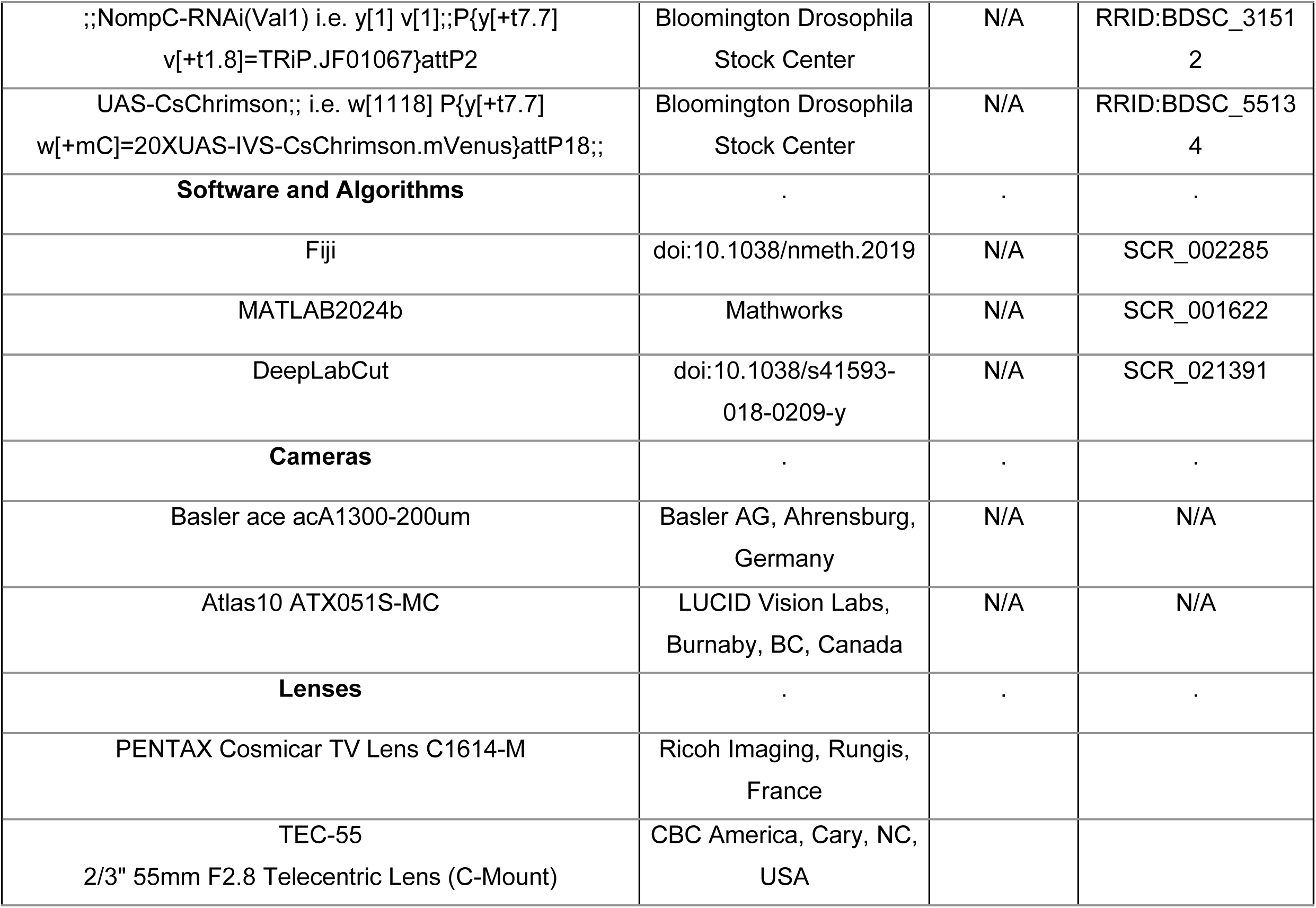
REAGENTS AND FLY STRAINS.

### Leg and wing preparations

Following CO_2 a_nesthesia, three to eight day-old flies were washed and sacrificed in ethanol. Dissection was performed in 0.1 mol l^-1^ PBS. Legs and wings were fixed in 4% paraformaldehyde (PFA) with glacial acetic acid for 45 min and washed with PBS. Wings and front, middle, and hind legs were collected separately to guarantee proper identification. No antibodies were needed due to strong endogenous fluorophore expression. Tissues were mounted in VECTASHIELD under a coverslip.

### Calcium imaging of leg muscles

We used five to eight days old male and female flies. eyg-Gal4 was used to drive the expression of CsChrimson ^76^ for CS activation, and MHC-LexA to drive the expression of GCaMP6f in leg muscles ^5,42^ on eyg>CsChrimson^muscle^ flies. GCaMP activity was monitored with a two-photon microscope (Bergamo II, Thorlabs). Flies carrying emptySplit-Gal4 instead of eyg-Gal4 were used as genetic controls (emptySplit>CsChrimson^muscle^).

We decapitated flies and sealed the neck with UV-curable glue and left on a moist petri dish to sort viable animals, indicated by the ability to right themselves and respond with grooming behavior when gently brushed. Flies were anesthetized on a chilled metal surface to remove wings and all legs except the left front leg, reducing optogenetic activation of other leg CS. Leg stumps were sealed with UV-curable glue to prevent hemolymph loss. Flies were glued parallel to a circular coverslip (Fig. 2, Fig. S3). Left front legs were gently glued to the coverslip to prevent motion artefacts during recordings. These were positioned between 30° to 140° femur-tibia joint angles and 120° to 190° tibia-tarsus joint angles.

The coverslip was fitted in the imaging chamber (CSC-25L, Bioscience Tools). The fly holder was then flipped so that the fly was underneath the coverslip on the opposite side to the objective. We added water on top of the coverslip and imaged with a water immersion objective (x20 NA 1.0 objective lens, Olympus XLUMPLFLN). CS were stimulated for 2s at 100Hz with a red light-emitting fiber (655 nm, 0.08 mW mm^-2^) pointed at the leg with a micromanipulator placed at roughly 1 cm distance. Stimulation was controlled and synchronized to the imaging session using the software ScanImage (MBF Bioscience).

Leg muscles were located using 1P wide-field fluorescence imaging (488 nm mounted LED; Thorlabs). Muscle activity was recorded via GCaMP6f signals (3Hz) under 2P imaging (920 nm Ti:Sapphire laser; MaiTai DeepSee, Newport Spectra-Physics) since it allowed for better muscle anatomical detail. Samples were stimulated at least four times per session, per leg segment.

### Behavioral experiments

One to two day-old flies were sorted under cold-anesthesia into vials with ATR. At three to five days-old eyg>GtACR1, eyg>GtACR1^brain^ and eyg>GFP flies were gently aspirated onto the setups (Fig. S8) and habituated for 10 to 20 minutes before experimentation. These setups are similar to previous studies ^16,58^ with modifications to the camera and software. The light-gated ion channel GtACR1 (*Guillardia theta* anion channelrhodopsin 1) was used to reversibly inhibit neurons via green light exposure (520 nm), hyperpolarizing them by transiently opening membrane channels permitting anion (chloride) influxes ^77^. GFP was used as a genetic control as it shares its genetic background w[*] and a similar base pair count as UAS-20xGtACR1.d.EYFP.

### Climbing assay

The chamber depicted in Supplementary Figure 8A was used for climbing assays. Vials contained five male and five female flies. An IR lamp (850 nm) with a sheet of diffuser foil was used for background lighting. A green-LED (520 nm) ring was mounted parallel to the vials and switched manually for optogenetic inhibition. A custom 3D-printed holder was used to support the upper section of the vials. We continuously recorded the vials with a monochrome camera (20 frames per second (FPS), 1280x1024 pixel resolution; Basler ace acA1300-200um, Basler AG, Ahrensburg, Germany). The two vials were simultaneously dropped from a height of approximately 10 cm, shaking the flies to the bottom and promoting climbing. Flies were allowed to climb for 10 s between drops. Rubbery foam on the holder’s bottom surface reduced the impact shock. Each pair of genotypes (eyg>GtACR1 *versus* eyg>GFP, eyg>GtACR1 *versus* eyg>GtACR1^brain^, eyg>GFP *versus* eyg>GtACR1^brain^) was dropped ten times per experiment. Five replicas were performed (50 flies per genotype). Control and inhibited trials were performed with the same flies, providing an internal control for the influence of the green light in climbing performance. We first ran the control paired trials (light-off) and, after a 10 min resting time, the same genotypes were tested for inhibition (light-on).

### Free-walking assay, high-level kinematics

Three flies were aspirated onto a 10 cm (diameter) inverted petri dish covered by a watch glass coated with SigmaCote to prevent ceiling walks (Fig. S8B, Fig. 3B, Fig. S5). We positioned two LED rings, one with green illumination (520 nm) for inhibition, another with IR illumination (830 nm) to record flies walking in the dark using a monochrome camera (20 FPS, 1280x1024 pixel resolution; Basler ace acA1300-200um, Basler AG, Ahrensburg, Germany) (Fig. S1B). A circular strip of diffusion foil was added in front of the IR LED to evenly distribute light. Trials consisted of 12 cycles (36 minutes total) of two minutes of darkness (control) and one minute of green light (inhibition) exposure.

### Free-walking assay, low-level kinematics

The setup (Fig. S8B) was as previously described in the “Free-walking, high-level kinematics” section, except for the dish and camera and is similar to previous studies ^16,58^. Single flies were placed onto a petri dish (diameter: 6 cm) enclosed by a watch glass coated with SigmaCote (Fig. 4-5, Fig. S6-7). We used a monochrome camera (200 FPS, 2000x2000 pixel resolution; Atlas10 ATX051S-MC, LUCID Vision Labs, Burnaby, BC, Canada). Single flies were recorded on a 200x200 pixels fly-centered cutout of the full camera resolution. Experiments consisted of cycles of two minutes in darkness (control) and one minute with green light (inhibition) for two hours.

### Side-view setup

A custom-made 3D printed chamber modified from a previous study ^57^ was used to record flies from the side (Fig. S8C, Fig. 6) consisting of: a top part housing green LEDs (520 nm) and a layer of diffusion foil to evenly distribute light on the walking corridor; two symmetrical side pieces, each housing two IR LEDs (730 nm) and a custom cut glass coated with SigmaCote to prevent climbing; one custom cut glass – the floor of the arena; a mirror, placed below the floor of the arena and oriented at 45° in relation to the camera and the floor; two custom cut glass pieces through which we filmed flies walking. We used a monochrome camera (20 FPS, 1280x1024 pixel resolution; Basler ace acA1300-200um, Basler AG, Ahrensburg, Germany) for video recording.

## DATA AND STATISTICAL ANALYSIS

### Calcium imaging

First, imaging data were baseline-corrected. For this, we manually selected a region of interest (ROI) for each respective session that was clearly separated from actual fluorescence and leg structures and that only contained general imaging noise. The activity of this background ROI was averaged and the resulting time course was subtracted from the activities of the muscle ROIs, thus correcting for general baseline fluctuations or potential brightness artifacts during stimulation. This was done for each session separately. Baseline calcium fluorescence (F_0)_ of inactive muscles was generally low and on a similar level as baseline fluorescence. This made the standard approach of reporting fluorescence changes relative to baseline (ΔF/F­_0)_ very sensitive to small changes in baseline and resulted in large inter -session differences in calculated ΔF/F­_0 v_alues between sessions. To make data more comparable and be able to pool all sessions, we therefore calculated the z-scores of peri-stimulus ROI signals that corresponded to sequences starting 2 s before stimulus onset and terminated 2 s after stimulus end (total duration 6 s). These standardized peri-stimulus signals were then averaged for a session and all session averages of a particular genotype were pooled.

### Climbing assay

To evaluate climbing performance we inspected the videos and manually annotated the times of all individual vial drops. These were defined as the onsets of each climbing trial. We then extracted the video frames 2.1 s after the drops, allowing a minimum climbing height to have a minimal agglomerate of pixels identifiable as one fly. We manually marked the outlines of the two vials in the extracted frames. The pixels in the two regions of interest (ROIs) were converted to binary black and white frames with an empirically determined threshold. We then calculated the climbed distance of all white pixels (flies) and pooled the data for all experiments, resulting in a distribution of climbed height per genotype for control and inhibited conditions. The non-parametric two-sided Wilcoxon signed-rank test was used to determine statistical significance.

### High-level kinematics: video extraction

We included walking trajectories above a speed threshold of 4 mm s^-1^ and below 30 mm s^-1^, excluding potential jumping instances. Walking bouts were termed “tracklets”. Each tracklet terminated when two flies intersected, spawning two new tracklets after separation. Tracklets were defined with a minimum walking duration of 0.5 s.

To extract bouts from each two hour experiment we: tracked the fly for experiment duration; segmented each experiment into manageable walking bouts; classified and sorted bouts regarding speeds, curvatures and distances to the center of the arena (to reconstruct the fly-centered 200x200 pixel cutout trajectory in a world-centered coordinate system). We focused on straight walking behavior (low curvature), encompassing walking bouts with a minimum length of 5 mm, and speeds ranging from 5 to 50 mm s^-1^, resulting in a selection of comparable walking bout segments (each with three to 20 step cycles). Segments were compared on a per-fly basis by selecting bouts with similar walking distances and speeds for the “light-off” and “light-on” conditions.

Speeds, trajectories and bout lengths were compared between flies walking in light-off *vs* light-on conditions. Non-optogenetic trials (RNAi genotypes) consisted of comparing the same genotypes, raised at either 18°C or 29°C (restrictive and permissive temperatures, respectively) walking for 36 minutes. Speeds from all tracklets were compiled into a matrix to calculate average speeds. The first 10 s of each light period and the first 30 s of each dark period were excluded, excluding potential startle responses and lingering inhibition effects during the “light-off” condition. The Kolmogorov–Smirnov test was used to calculate statistical significance. A p-value < 0.05 was considered statistically significant. Due to a high “n” (often n > 10,000) the Kolmogorov–Smirnov statistic (*D*) was used to indicate effect size (Supplementary Table 1). The *D* value represents the maximum difference between cumulative distributions.

### Low-level kinematics

Walking speed influences leg stepping kinematics as well as interleg coordination ^15,57^. Therefore, we speed-matched each “light-on” video to a control video (“light-off”) whenever the behavioral phenotype allowed it, thus minimizing the effect speed has on kinematics (e.g.: eyg>GtACR1^brain^). Using a light-off *vs* light-on protocol allowed us internal control comparisons even when the walking speeds were drastically different due to the elicited phenotypes, e.g. eyg>GtACR1. Selected videos were automatically annotated via DeepLabCut (DLC) ^16,58,78^ trained on previously recorded and manually annotated walking trials performed by a wild type strain (Berlin-K). DLC-based annotation errors were corrected manually.

For each leg, swing and stance phases were defined via the speed of the tarsal tips relative to the ground. Speeds close to zero thereby corresponded to the stance phase. For all further analysis steps, kinematic data were transformed into a fly-centric coordinate system, based on the vector between neck and abdomen. Lift-off and touchdown times were defined by the transition to the swing phase and stance phase, respectively. The tarsal positions at these times were defined as the posterior extreme positions (PEPs at lift-off) and anterior extreme positions (AEPs at touchdown). Swing duration was defined as the time between lift-off and touchdown; stance duration was defined as the time between touchdown and liftoff. The duration of a complete step was defined as the time between two consecutive lift-offs. Stance trajectories were defined as the trajectory of tarsal tips between an AEP and the subsequent PEP; stance amplitudes were defined as the distance between this AEP and PEP. Phase relationships were defined calculated between adjacent legs (ipsilaterally and contralaterally),

Individual kinematic parameters were normalized to the arithmetic mean of the control condition for “light-on” comparisons. Normalization allowed for direct comparison between flies and pooling of all data points as kinematic parameters naturally differ between flies. The non-parametric two-sided Wilcoxon signed-rank test was used for kinematic parameters. For phase relationships, we used a non-parametric variant of the Kruskal–Wallis test for circular statistics ^79^. Due to the high sample size, many comparisons are statistically significant. We therefore focus on actual effect sizes.

### Side-view

Fly ground clearance was analyzed over an interval spanning 1.5s before and after each light event (i.e. light onset or offset). Valid events were selected using a custom-written graphical user interface to include only behavioral sequences in which the animals walked predominantly straight at a relatively constant speed, did not touch or climb the recording chamber walls, and remained stationary during standing prior to the light event. The contour of the fly’s body was detected in each frame using a threshold operation, followed by fitting an ellipse to the resulting binary blob using the OpenCV framework (version 4.10.0) ^80^. The major and minor axes of the fitted ellipse corresponded to the anterior-posterior and ventral-dorsal axes of the body, respectively. Ground clearance was defined as the vertical distance from the chamber floor to the fly’s notum, as approximated by the upper endpoint of the ellipse’s minor axis. To account for inter-individual differences in body size and the step-cycle-related vertical motion inherent to walking, we computed relative ground clearance for each time point of a sequence by normalizing to the mean height value of the pre-event period (30 frames; 1.5 seconds) and smoothing the resulting time series using a Gaussian filter (σ = 1.75). For statistical analysis, we computed the median relative height and its 95% confidence interval using the bias-corrected and accelerated bootstrap method.

## Acknowledgements

We thank Salil S. Bidaye and the Max Planck Florida Institute for hosting RDC during the calcium imaging two-photon microscopy experiments. We thank Nino Mancini for the teaching and technical assistance during the stay at Salil S. Bidaye’s lab. We thank Mehrdad Ghanbari, Sima Seyed-Nejadi and Sherylane Seeliger for technical assistance. We thank the Imaging Facility in the Biocenter Cologne and its support staff.

## Author contributions

- RDC, EAG, TB and AB conceived and designed the research.
- RDC, MD, TB and AB developed the behavioral setups.
- RDC, EAG and AP performed experiments and analyzed data.
- MH, VG and TB developed scripts and software for behavioral recordings and data analysis.
- RDC, EAG, MH, TB and AB interpreted the results.
- RDC, AP, MH and TB designed and prepared figures.
- RDC and AB drafted the manuscript.
- RDC, EAG, MH, TB and AB edited and revised the manuscript. All authors approved the submitted version.

## Supplementary figures and video legends

### Supplementary video1

**Supplementary Figure 1:**
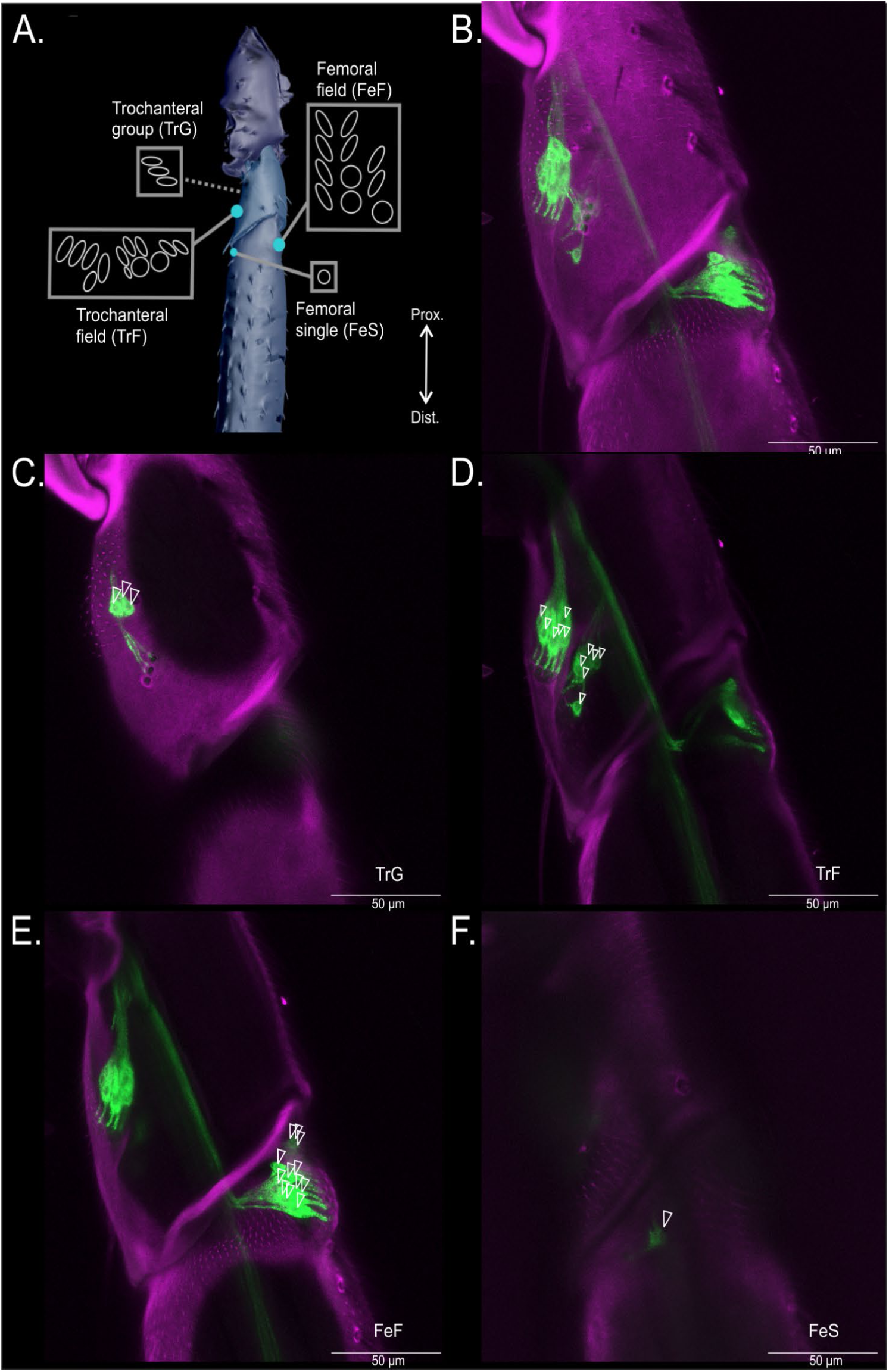
**A)** Photography taken with a Sony Alpha 7 camera mounted over a fluorescent lamp. The image depicts a male *D.mel* eyg-Gal4 fly expressing GFP (N= 2 male, N =1 female). eyg>GFP shows labeling on all leg CS, as well as those present in the halteres and wings. Proboscis and antennae also show labeling, possibly due to presence of mechanotransductors in these structures. **B)** Max intensity projection of eyg-positive cells in a front leg (10x) of a female eyg>GFP,RedStinger *D.mel* (n=3). Cyan: background autofluorescence of leg cuticle (666-800nm). Green: GFP signal from eyg-Gal4 labeled cells (502-517nm), replicating the locations of the SEM-based scheme in Fig. 1B. Red: dsRed signal (575-590nm) labeling the nuclei of eyg positive cells (UAS-RedStinger). Yellow: summation of the wavelengths of the green and red channels, showing the coincidence of CS and their nuclei respectively. **C)** Max intensity projection a tile scan of eyg-positive cells in a front leg’s tarsa (40x) of a female eyg>GFP,RedStinger *D.mel* (n=3). Description of channel intervals as in Fig. S1A. **D)** Max intensity projection of eyg-positive cells in a front leg’s tibia and tarsa (40x) of a female eyg>GFP,RedStinger *D.mel* (n=3). Description of channel intervals as in Fig. S1A. **E)** Max intensity projection of eyg-positive cells in a front leg’s coxa and trochanter and tibia (40x) of a female *D.mel* eyg>GFP,RedStinger (n=3). Description of channel intervals as in Fig. S1A. **F)** Max intensity projection a tile scan of eyg-positive cells in a wing (10x) of a female eyg>GFP,RedStinger *D.mel* (n=3). Description of channel intervals as in Fig. S1A.

**Supplementary Figure 2:**
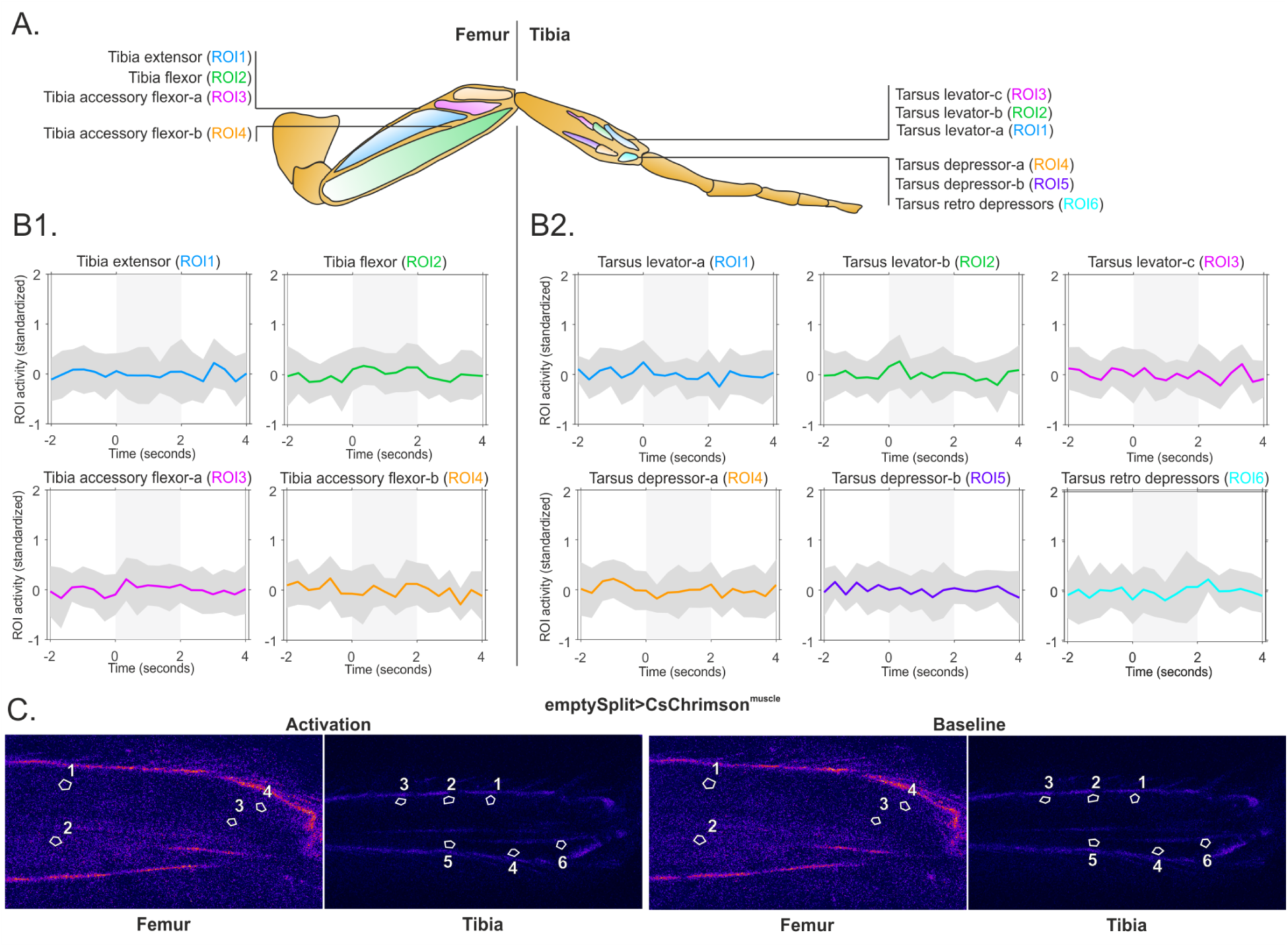
**A)** Illustration based on ^27^ depicting the distribution of CS in the trochanteral and femoral leg segments. Left to right, top to bottom: TrG: trochanteral group; TrF: trochanteral field; FeF: femoral field; FeS: femoral single CS. **B)** Full stack of a Max intensity projection of eyg-positive cells in a front leg’s trochanter and tibia (63x) of a female *D.mel* (n=3). Magenta: background autofluorescence of leg cuticle (666-800nm). Green: GFP signal from eyg-Gal4 labeled cells (502-517nm). **C)** Max intensity projection of a selection of 34 images from the full stack in B) showing three eyg-positive cells in a front leg’s trochanteral group (TrG) (63x) of a female *D.mel*. Description of channel intervals as in Fig. S2B. **D)** Max intensity projection of a selection of 71 images from the full stack in B) showing twelve eyg-positive cells in a front leg’s trochanteral field (TrF) (63x) of a female *D.mel*. Description of channel intervals as in Fig. S2B. **E)** Max intensity projection of a selection of 111 images from the full stack in B) showing twelve eyg-positive cells in a front leg’s femoral field (FeF) (63x) of a female *D.mel*. Description of channel intervals as in Fig. S2B. **F)** Max intensity projection of a selection of 24 images from the full stack in B) showing one eyg-positive cell (FeS) in a front leg’s femur (63x) of a female *D.mel*. Description of channel intervals as in Fig. S2B.

- **1)** Video of the z-stack of eyg-Gal4 positive cells in a front leg’s trochanter and tibia (63x) of a female *D.mel* (N=3). Total number of eyg-positive cell somas are displayed and numbered. Order of appearance reflects the depth sequence. Magenta: background autofluorescence of leg cuticle (666-800nm). Green: GFP signal from eyg-Gal4 labeled cells (502-517nm).

**Figure S3:**
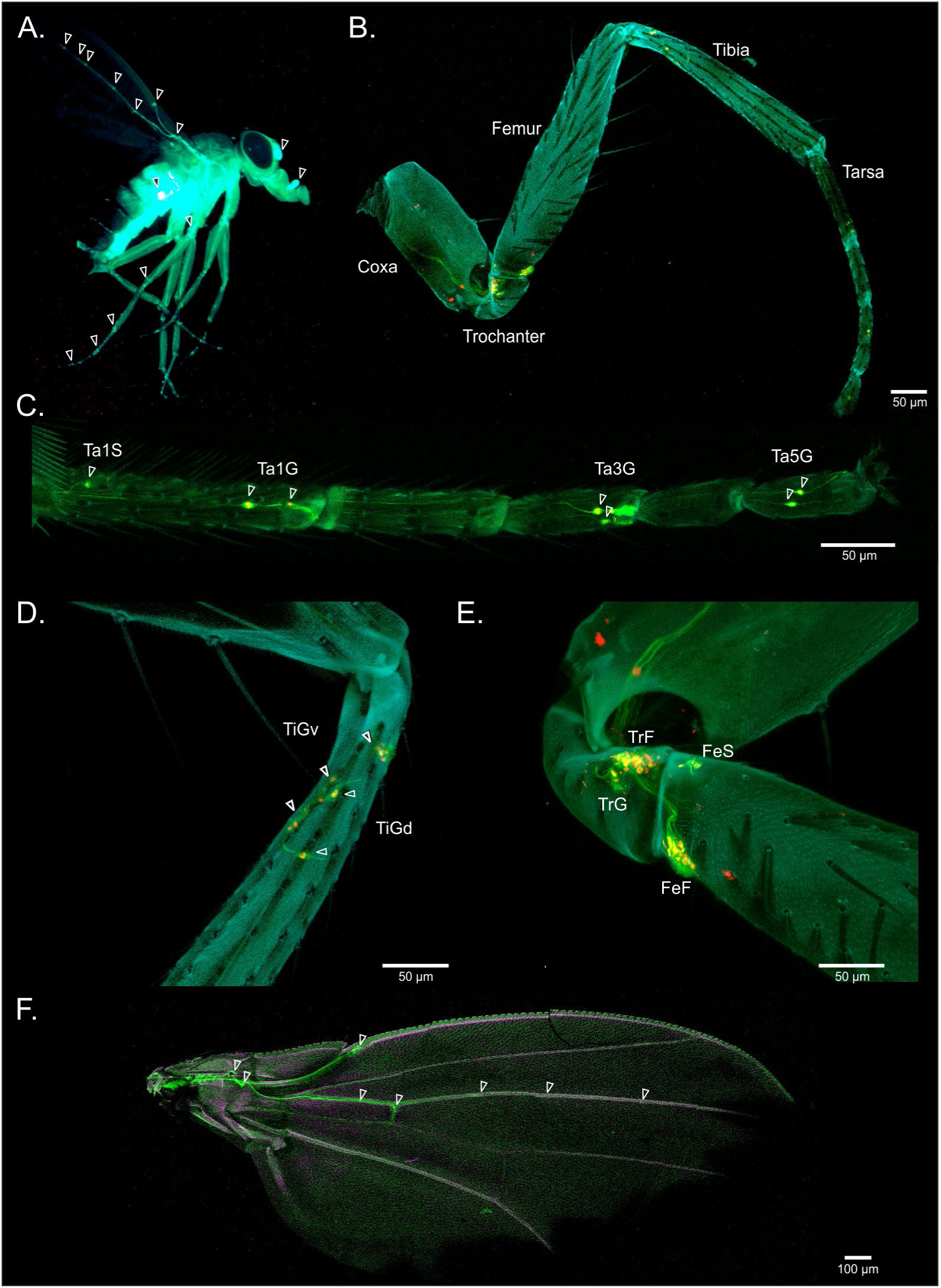
**A)** Depiction of imaged femoral and tibial leg muscles where numbers indicate muscle ROIs in panels B and C. For all muscle nomenclatures please see Azevedo *et al.* 2024 (supplementary materials) ^45^ and Soller *et al.* 2004 ^44^. **B) B1)** Quantification of femoral muscle activity events in emptySplit>CsChrimson^muscle^ flies (y-axis: standardized activity within ROI) elicited by a 2 s red light pulse (x-axis: prestimulus (-2 to 0 seconds), stimulus (0 to 2 seconds) and poststimulus (2 to 4 seconds) intervals. ROIs 1 to 4 correspond to leg muscles in graph titles. Colored line depicts mean standardized activity, grey shade depicts SD of activity from different preparations (N=14, pooled data from extended (N=3, n=4), neutral (n=4) and flexed (N=5, n=6) leg positions). **B2)** Quantification of tibial muscle activity events in emptySplit>CsChrimson^muscle^ flies (y-axis: standardized activity within ROI) elicited by a 2 s red light pulse (x-axis: prestimulus (-2 to 0 seconds), stimulus (0 to 2 seconds) and poststimulus (2 to 4 seconds) intervals. ROIs 1 to 6 correspond to leg muscles in graph titles. Colored line depicts mean standardized activity, grey shade depicts SD of activity from different preparations (N=12, pooled data from extended (N=3), neutral (N=5) and flexed (N=4) leg positions). **C)** Depiction of a single muscle activity event (left, “Activation”) and baseline muscle activity (right, “Baseline”) within the femoral or tibial leg segments of the same emptySplit>CsChrimson^muscle^ fly. ROI numbering corresponds to muscles analyzed in (B).

### Supplementary videos 2-5

- **2)** Depiction of muscle activity events for the femoral segment of an eyg>CsChrimson^muscle^ fly’s leg in a neutral position. Events represent intramuscular calcium bursts correlated with activity. Colored bar represents pulse of red light (2 seconds) acquired at 3 Hz. The video is playing at 15 FPS. A Gaussian blur was applied (0.3).
- **3)** Depiction of muscle activity events for the tibial segment of an eyg>CsChrimson^muscle^ fly’s leg in a neutral position. Events represent intramuscular calcium bursts correlated with activity. Colored bar represents pulse of red light (2 seconds) acquired at 3 Hz. The video is playing at 15 FPS. A Gaussian blur was applied (0.3).
- **4)** Depiction of muscle activity events for the femoral segment of an emptySplit>CsChrimson^muscle^ fly’s leg in a neutral position. Events represent intramuscular calcium bursts correlated with activity. Colored bar represents pulse of red light (2 seconds) acquired at 3 Hz. The video is playing at 15 FPS. A Gaussian blur was applied (0.3).
- **5)** Depiction of muscle activity events for the tibial segment of an emptySplit>CsChrimson^muscle^ fly’s leg in a neutral position. Events represent intramuscular calcium bursts correlated with activity. Colored bar represents pulse of red light (2 seconds) acquired at 3 Hz. The video is playing at 15 FPS. A Gaussian blur was applied (0.3).

**Supplementary Figure 4:**
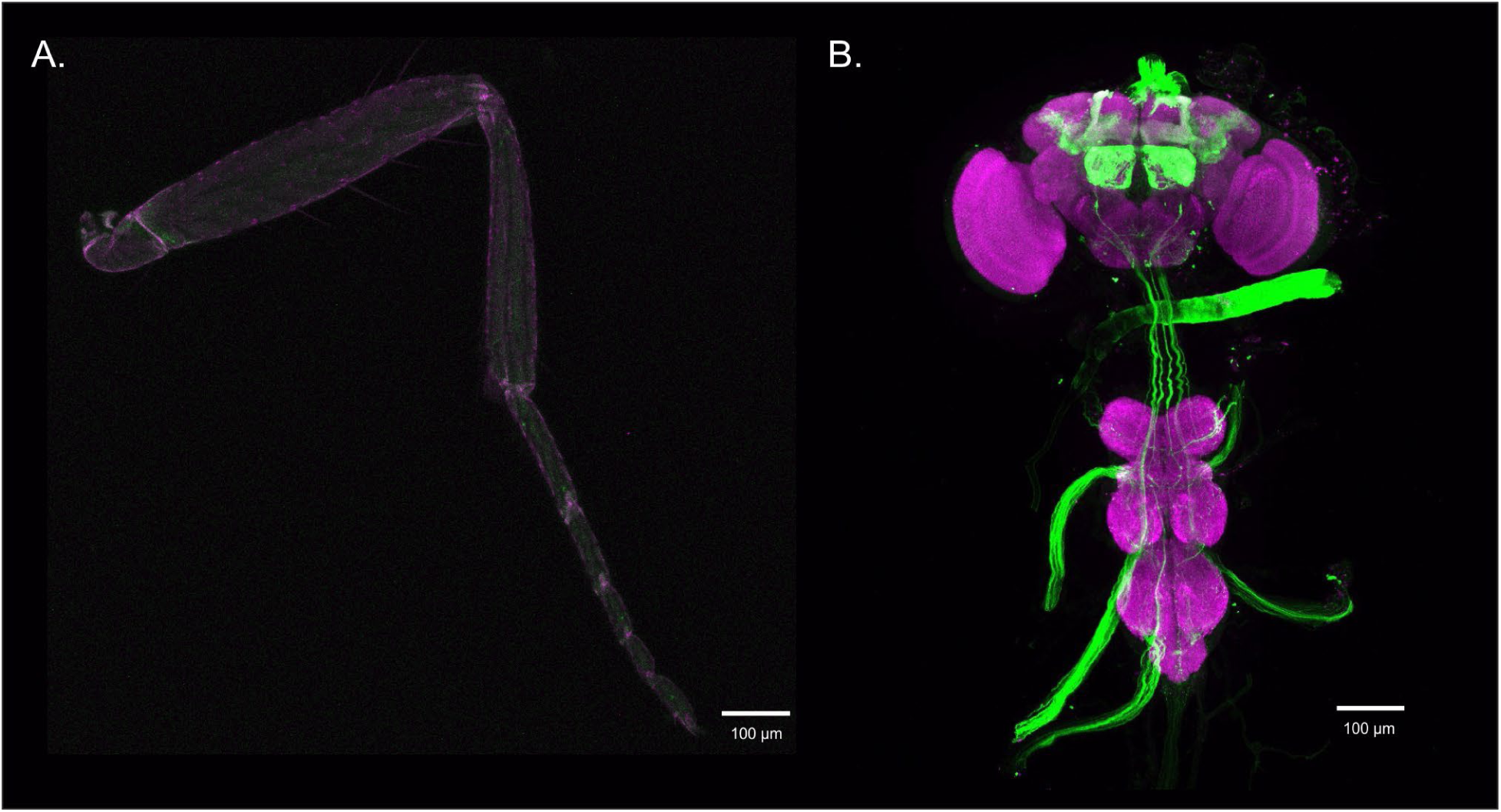
**A)** Max intensity stack of a eyg>GtACR1^brain^ *D.mel*’s middle leg (10x) expressing EYFP in eyg-positive neurons, under the restriction of the brain labeling construct. Magenta: background autofluorescence of CNS (666-800nm). Green/white (post-processing editing for increased contrast): GFP signal from eyg-Gal4 labeled cells (502-517 nm) (N=3). **B)** Max intensity stack of a eyg>GtACR1^brain^ *D.mel* CNS (10x) expressing EYFP in eyg-positive neurons, under the restriction of the brain labeling construct. Magenta: nc82 signal of CNS labeled with Alexa 405 (398-412nm). Green/white (post-processing editing for increased contrast): GFP signal from eyg-Gal4 labeled cells (502-517 nm) (N=3).

### Supplementary videos 6-11

- **6)** Depiction of a climbing assay experiment comparing eyg>GtACR1 (left vial) to eyg>GFP (right vial) in control conditions (in the dark).
- **7)** Depiction of a climbing assay experiment comparing eyg>GtACR1 (left vial) to eyg>GFP (right vial) in the optogenetic inhibition condition (green light on condition).
- **8)** Depiction of a climbing assay experiment comparing eyg>GtACR1 (left vial) to eyg>GtACR1^brain^ (right vial) in control conditions (in the dark).
- **9)** Depiction of a climbing assay experiment comparing eyg>GtACR1 (left vial) to eyg>GtACR1^brain^ (right vial) in the optogenetic inhibition condition (green light on condition).
- **10)** Depiction of a climbing assay experiment comparing eyg>GtACR1^brain^ (left vial) to eyg>GFP (right vial) in control conditions (in the dark).
- **11)** Depiction of a climbing assay experiment comparing eyg>GtACR1^brain^ (left vial) to eyg>GFP (right vial) in the optogenetic inhibition condition (green light on condition).

**Supplementary Figure 5:**
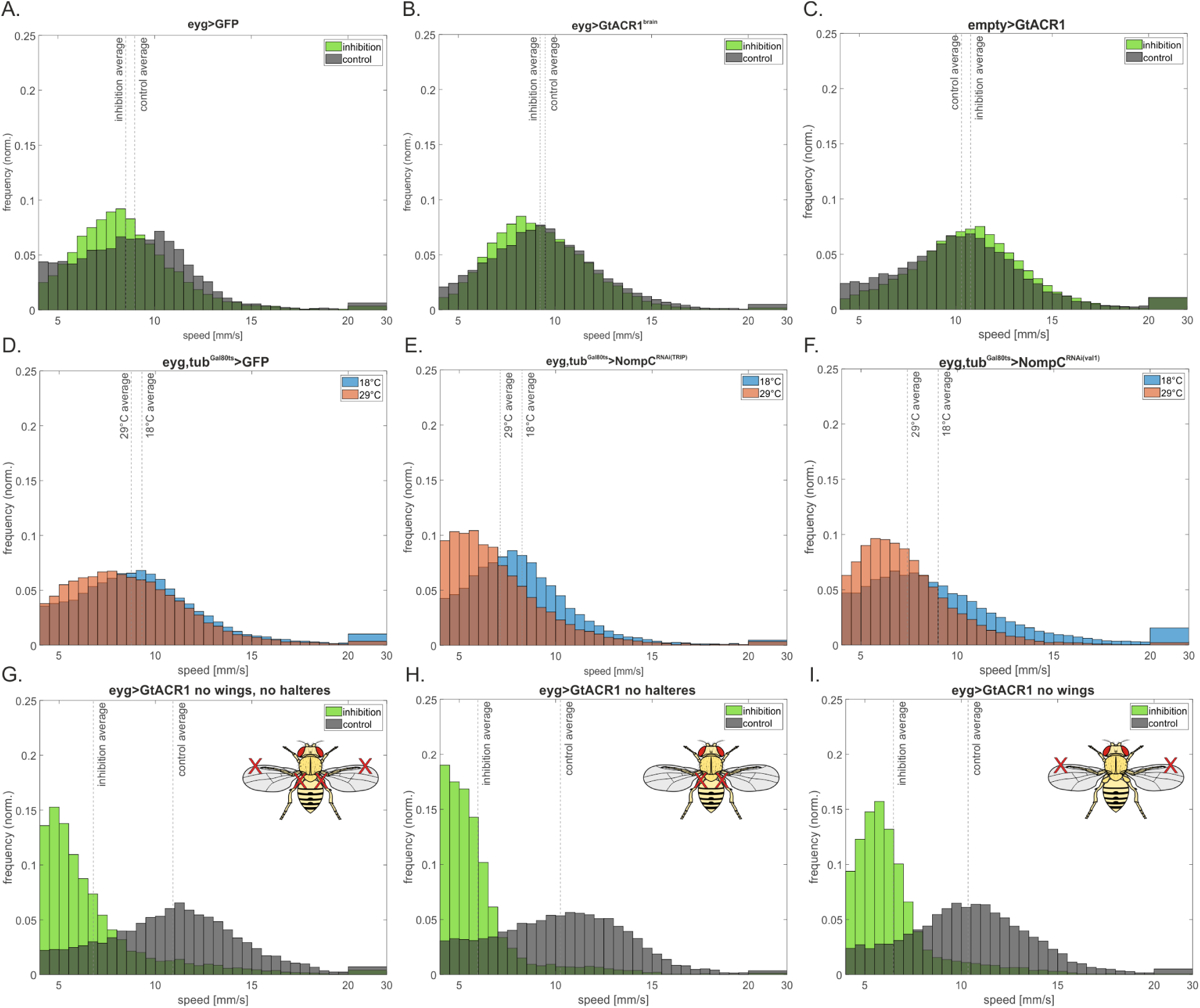
**A)** Frequency of speed distribution from walking bouts of the eyg>GFP negative genetic control flies (N=30). Description of graphic features in Fig. 3B legend. **B)** Frequency of speed distribution from walking bouts of the eyg>GtACR1^brain^ brain restricting control (N=27). Description of graphic features in Fig. 3B legend. **C)** Frequency of speed distribution from walking bouts of the empty>GtACR1 negative genetic control for the GtACR1-positive UAS promoter line, expressed under the empty tissue driver empty-Gal4, instead of the CS driver eyg-Gal4 (N=30). **D)** Frequency of speed distribution from walking bouts of the temperature control eyg,tub^Gal80ts^>GFP for the NompC-RNAi UAS promoter line, expressing GFP under the CS driver eyg-Gal4. Flies were maintained at 18°C (blue, N=36) or 29°C (red: N=42) post eclosion. Description of graphic features in Fig. 3B legend. **E)** Frequency of speed distribution from walking bouts of the positive genetic control eyg,tub^Gal80ts^>NompC^RNAi(TRIP)^. RNA translation was restricted (Blue: 18°C, N=18) or permitted (Red: 29°C, N=18) via the tubulin-Gal80^ts^ temperature sensitive construct. Flies grown at the permissive temperature of 29°C express an interference RNA (NompC-TRIP-RNAi) for the mechanoreceptor NompC in eyg-positive cells. Description of graphic features in Fig. 3B legend. **F)** Frequency of speed distribution from walking bouts of the positive genetic control eyg,tub^Gal80ts^>NompC^RNAi(val1)^. RNA translation was restricted (Blue: 18°C, N=18) or permitted (Red: 29°C, N=18) via the tubulin-Gal80^ts^ temperature sensitive construct. Flies grown at the permissive temperature of 29°C express an interference RNA (NompC-valium1 insertion-RNAi) for the mechanoreceptor NompC in eyg-positive cells. Description of graphic features in Fig. 3B legend. **G-I)**Controls for walking performance of eyg>GtACR1 in the absence of proprioceptive structures (amputated wings and halteres (Fig. S4D.; N=24), halteres (Fig. S4E.; N=18) and wings (Fig. S4F; N=24)). These controls (grey) were also tested under optogenetic inhibition (green) to solidify the leg-CS labeling phenotype (Fig. 5A). Description of graphic features in Fig. 5A legend. *Drosophila melanogaster* pictures are based on public domain drawing by Madboy74, CC0, via Wikimedia Commons.

**SUPPLEMENTARY TABLE 1:** Kolmogorov–Smirnov statistical test for walking speed distribution upon CS inhibition in *Drosophila*.

| *Drosophila* genotype | Kolmogorov–Smirnov D value for walking speed distribution |
| --- | --- |
| *eyg>GtACR1* | 0.4881 |
| *eyg>GFP* | 0.1272 |
| *eyg>GtACR1<sup>brain</sup>* | 0.0474 |
| *eyg,tub<sup>Gal80ts</sup>>NompC<sup>RNAi(TRIP)</sup>* | 0.2357 |
| *empty>GtACR1* | 0.0750 |
| *eyg,tub<sup>Gal80ts</sup>>NompC<sup>RNAi(val1)</sup>* | 0.2152 |
| *eyg,tub<sup>Gal80ts</sup>>GFP* | 0.0817 |
| *eyg>GtACR1* no halteres, no wings | 0.5829 |
| *eyg>GtACR1* no halteres | 0.6480 |
| *eyg>GtACR1* no wings | 0.6256 |

### Supplementary videos 12 and 13

- **12)** Example video of a eyg>GtACR1 trial in control (dark) condition. Colored lines represent swing and stance phases color coded as Fig. 6B2.
- **13)** Example video of a eyg>GtACR1 trial in inhibited (light on) condition. Colored lines represent swing and stance phases color coded as Fig. 6B2.

**Supplementary Figure 6:**
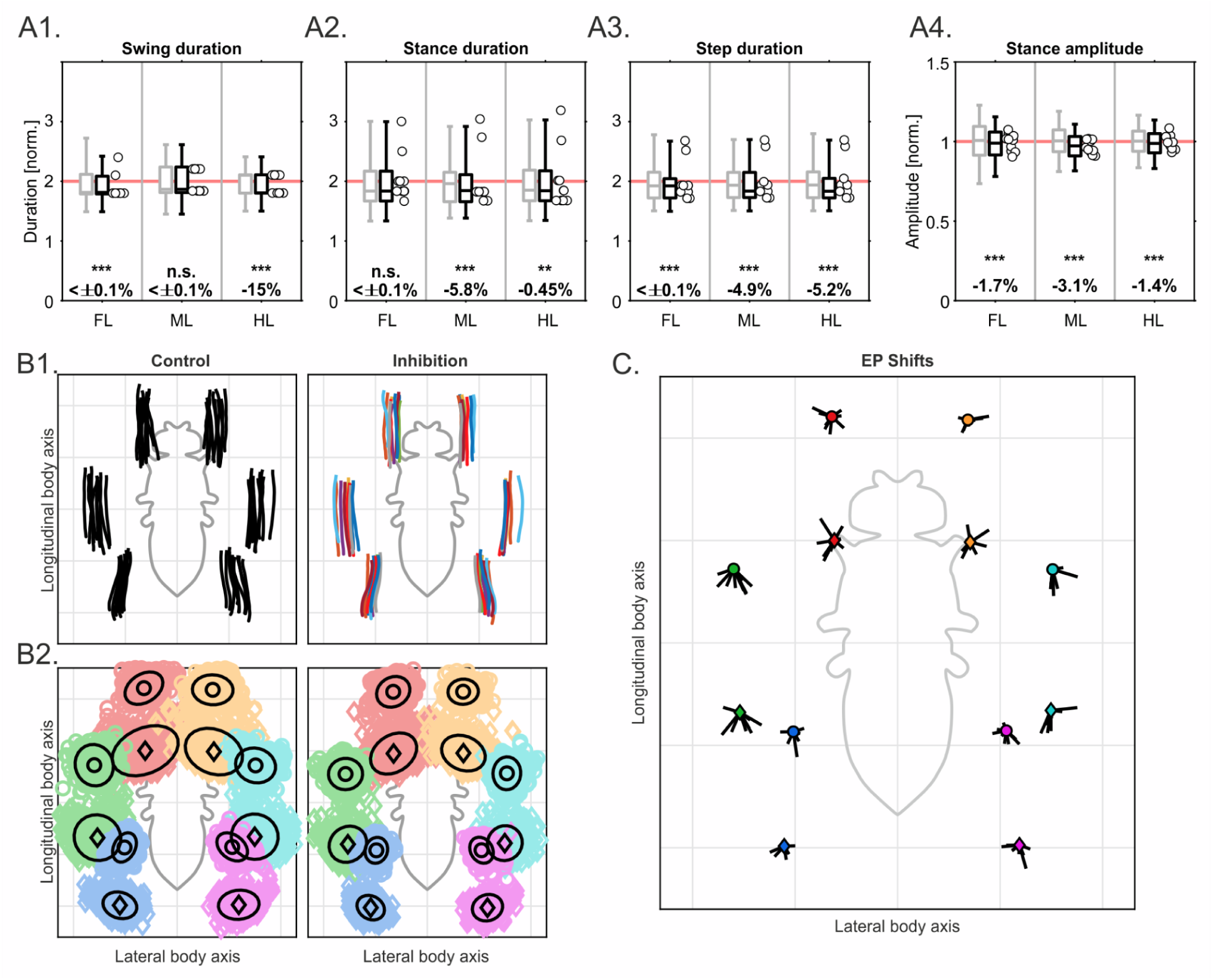
**A)** A1-4) Normalized swing, stance and step durations and stance amplitudes produced during straight walking bouts (minimum 5 steps per leg) by eyg>GtACR1^brain^ flies (y-axis). Grey and black boxplots represent walking bouts produced by the same fly in the dark and under green light. White dots: individual average of each inhibited fly. Steps from front (FL), middle (ML) and hind legs (HL) were paired (x-axis). Normalization was done on a per fly basis to their mean values in the control condition (indicated as 1, light red reference line). Box plots (whiskers indicate 1% and 99% percentiles) are combined normalized data for all flies; circles indicate medians for individual flies. Combined data in the inhibited condition were tested against a median of 1 with a two-sided Wilcoxon signed rank test. Significance levels of these tests are given as non-significant (n.s.), *P≤0.05, **P≤0.01 and ***P≤0.001. Effect sizes are given as the difference between the median in the inhibited condition and 1 (normalized control condition). Each fly (N=10 has 10 control and 10 inhibition videos (n=200). **B) B1)** Average stance trajectories for each individual fly shown in fly-centered coordinate space for the control (left: black lines) and inhibited condition (right: lines color coded per individual). **B2)** distribution of anterior (AEPs: colored circles) and posterior extreme positions (PEPs: colored diamonds) with their mean positions (black circles and diamonds) and 95% percentiles (black ellipses). Colors represent the front (right: red; left: yellow), middle (right: green; left: cyan) and hind legs (right: blue; left: pink). **C)** Extreme position (EP) shifts. For each leg, each vector (black lines) corresponds to one fly and displays the average shift of AEPs (top) and PEPs (bottom) in the inhibited condition, compared with their respective mean positions in the control condition (circles, color coded as B2.).

### Supplementary videos 14 and 15

- **14)** Example video of a eyg>GtACR1^brain^ trial in control (dark) condition. Colored lines represent swing and stance phases color coded as Fig. 6B2.
- **15)** Example video of a eyg>GtACR1^brain^ trial in inhibited (light on) condition. Colored lines represent swing and stance phases color coded as Fig. 6B2.

**Supplementary Figure 7:**
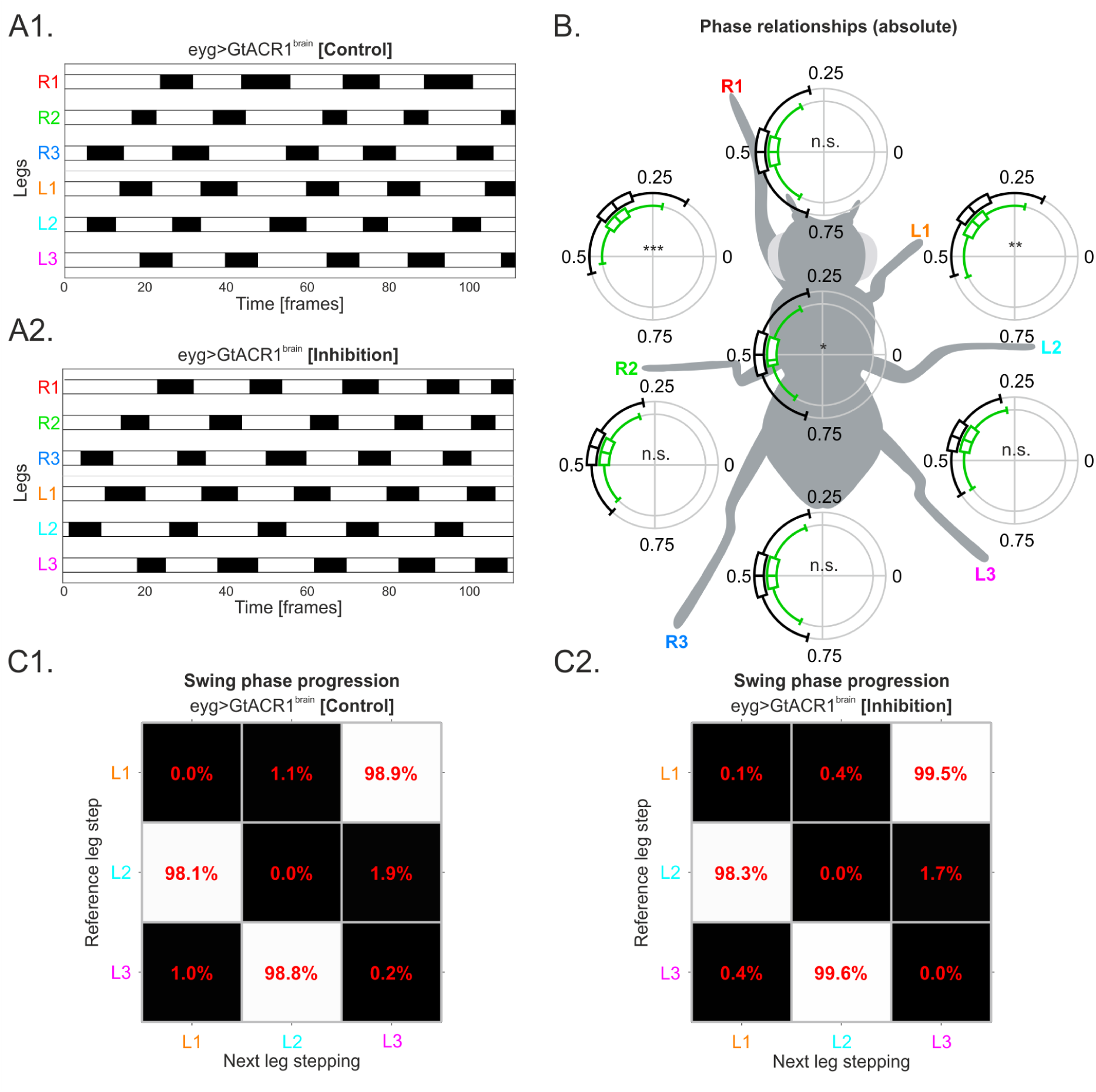
**A)** A1) Footfall patterns of example trial from a eyg>GtACR1^brain^ fly walking in the dark (control) and **A2)** light (inhibition) performing a straight walking bout covering approximately the same distance. The x-axis represents time in frames. In the y-axis R1, R2, R3 and L1, L2, L3 respectively account for the right and left front, middle and hind legs. White and black bars respectively represent the legs stance and swing phases. **B)** Absolute phase relationships for eyg>GtACR1^brain^ flies. Description of graphic features in Fig. 8B. legend. Each fly (N=10) has 10 control and 10 inhibition videos (n=200). **C)** Depiction of back-to-front ^58,59^ swing phase progression between ipsilateral left legs for control **C1)** and inhibited **C2)** walking bouts of eyg>GtACR1^brain^ flies. Description of graphic features in Fig. 8C. legend. Each fly (N=10) has 10 control and 10 inhibition videos (n=200).

### Supplementary videos 16 to 23

- **16)** Example video of a walking eyg>GtACR1 fly filmed from the side depicting its ground clearance change upon light onset (inhibition). Purple marker represents the notum position for each frame, blue markers represent the notum position at each past frame. Square at the upper right corner represents light status: off, empty square; on filled square.
- **17)** Example video of a walking eyg>GtACR1 fly filmed from the side depicting its ground clearance change upon light offset (cessation of inhibition). Description of graphic features in Supplementary video 16 legend.
- **18)** Example video of a walking eyg>GFP fly filmed from the side depicting its ground clearance change upon light onset (inhibition). Description of graphic features in Supplementary video 16 legend.
- **19)** Example video of a walking eyg>GFP fly filmed from the side depicting its ground clearance change upon light offset (cessation of inhibition). Description of graphic features in Supplementary video 16 legend.
- **20)** Example video of a standing eyg>GtACR1 fly filmed from the side depicting its ground clearance change upon light onset (inhibition). Description of graphic features in Supplementary video 16 legend.
- **21)** Example video of a standing eyg>GtACR1 fly filmed from the side depicting its ground clearance change upon light offset (cessation of inhibition). Description of graphic features in Supplementary video 16 legend.
- **22)** Example video of a standing eyg>GFP fly filmed from the side depicting its ground clearance change upon light onset (inhibition). Description of graphic features in Supplementary video 16 legend.
- **23)** Example video of a standing eyg>GFP fly filmed from the side depicting its ground clearance change upon light offset (cessation of inhibition). Description of graphic features in Supplementary video 16 legend.

**Supplementary Figure 8:**
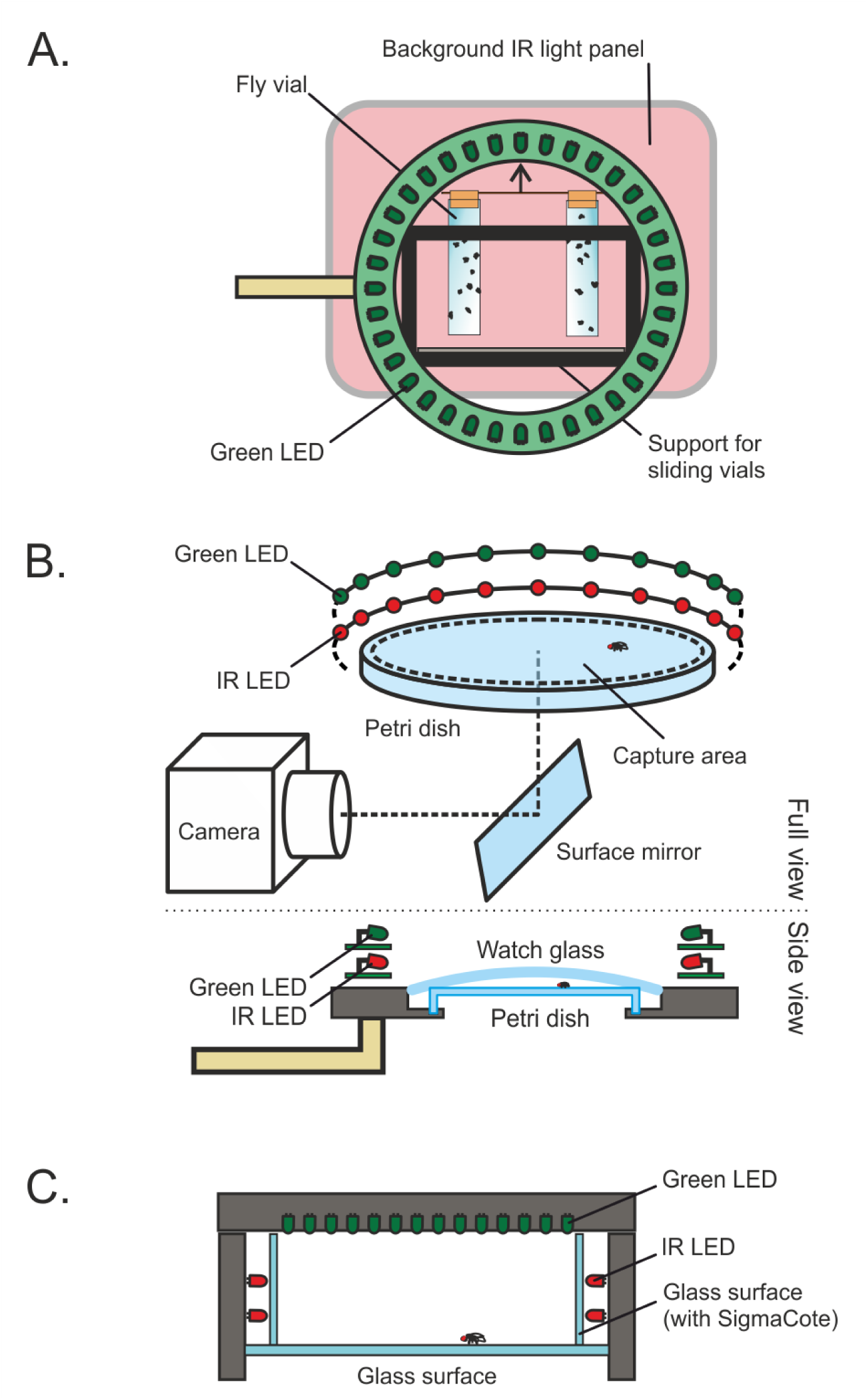
**A)** Climbing assay setup. The camera was located from the reader’s perspective. A ring with green LEDs shone on the vials, illuminated from the background using an infrared light panel. Vials were pulled up in the direction of the arrow and dropped, sliding in the vial support and cushioned at the bottom when hitting the spongy ground surface. **B)** Free walking setup. Full view: flies walked on a petri dish (diameter: 10 cm for high-level kinematics; 6 cm for low-kinematics) and IR camera captured all walking bouts performed on the capture area though the surface mirror reflection. IR LEDs allowed for camera detection and optogenetics were applied via green LEDs. Side view: flies were enclosed between the petri dish and a watch glass with SigmaCote on the inside layer, preventing ceiling walks. **C)** Side view setup. The camera was filming from the perspective of the reader. Glass surfaces enclosed the fly from all sides, except the top. Vertical surfaces were coated with SigmaCote to prevent climbing. Four IR LEDs illuminated the scene and green LEDs on top lid were used for optogenetic inhibition.

